# α-Synuclein aggregates induce mitochondrial damage and trigger innate immunity to drive neuron–microglia communication

**DOI:** 10.1101/2025.06.23.661105

**Authors:** Ranabir Chakraborty, Stephanie Maya, Veronica Testa, Jara Montero-Muñoz, Takashi Nonaka, Masato Hasegawa, Antonella Consiglio, Chiara Zurzolo

## Abstract

Tunneling nanotubes (TNTs) enable direct intercellular transfer of macromolecules, organelles, and pathogenic protein aggregates. While α-synuclein (α-Syn) aggregates are known to promote TNT formation, the underlying mechanisms remain poorly defined. Here, using human neuronal and microglial cell lines, as well as iPSC-derived dopaminergic neurons and microglia, we show that α-Syn aggregates induce severe mitochondrial damage, leading to cytosolic release of mitochondrial DNA (mtDNA) and activation of the cGAS–STING–NF- κB–IRF3 pathway. This innate immune response drives actin cytoskeleton remodeling and the formation of TNT-like structures, promoting intercellular transfer of α-Syn from neurons to microglia. Additionally, neuronal cells transfer damaged mitochondria to microglia, where they undergo lysosome-mediated degradation. Neuron-to-microglia communication under α-Syn-induced stress also triggers a bystander inflammatory response in microglia, suggesting a neuroimmune activation. Our findings identify mitochondrial damage and STING-mediated inflammation as key drivers of TNT formation and α-Syn propagation, highlighting new potential targets to modulate disease progression in Synucleinopathies.

## Main

Aggregation of α-Synuclein (α-Syn) is commonly observed in Synucleinopathies such as Parkinson’s (PD), dementia with Lewy bodies, and multiple systems atrophy^1–4^. Physiologically a cytosolic protein existing as a monomer, α-Syn has been reported to have association affinities to curved membranes such as synaptic vesicles, thereby implicating them in regulating vesicular dynamics at pre-synaptic terminals^5,6^. Mutations or duplication/triplication of the gene encoding α-Syn (*SNCA*) lead to pathological manifestations pertaining to aggregation of the protein^7^, characterized by significant loss of dopaminergic neurons in the Substantia Nigra pars compacta (SNpc).

Besides *SNCA*, several other genes have also been implicated in the pathogenesis of familial PD such as *PARKIN*, *PINK-1*, and *DJ-1*, protein products of which are essential for mitochondrial homeostasis^8–10^. Consequently, the pathogenesis of PD has been associated with mitochondrial dysfunction^11,12^. Interestingly, α-Syn has also been reported to interact with mitochondrial membrane, and irreversibly translocate to the matrix in a concentration-dependent manner, leading to impairment in ATP production^13,14^. Functional compromise of mitochondria also occurs as a result of oligomeric α-Syn binding with high affinity to the translocase of outer mitochondrial membrane TOMM20, causing impaired protein import into the organelle necessary to sustain physiological functions^15^. Significant damage to mitochondria leads to a loss of membrane integrity, allowing for mitochondrial ligands to be released into the cytoplasm, referred to as damage-associated molecular patterns (DAMPs)^16^. Among the different mitochondria-derived DAMPs is mitochondrial DNA (mtDNA) which share similarities with bacterial DNA owing to the endosymbiotic origin of these organelles^17,18^. When released in the cytosol, mtDNA triggers an innate immune response via engagement of cytosolic nucleic acid sensors such as cyclic GMP-AMP synthase (cGAS), leading to robust neuroinflammation^19^. Such inflammatory responses involving both pro-inflammatory cytokines and type I interferons are dependent on the activation of stimulator of interferon genes (STING), a phenotype observed in several neurodegenerative diseases like Alzheimer’s^20,21^, Parkinson’s^22,23^, Huntington’s^24^, and amyotrophic lateral sclerosis^25^.

A common pathological hallmark of neurodegenerative pathologies is their progressive spread to different regions of the brain. *In vitro*, tunneling nanotubes (TNTs) have emerged as a major route of aggregates such as α-Syn to directly spread not only between neurons^26,27^, but also between neurons and glia^28–30^. Since its initial demonstration in 2004^31^, TNTs are now recognized as unique facilitators of specialized communication between connected cells, also allowing for exchange of functional organelles such as mitochondria to rescue diseased phenotypes of cells^32–35^. Given the specialized nature of these structures which can be up- regulated “on-demand”, the field has a general consensus that stressors of different natures can promote TNT-mediated intercellular communication^36,37^. Importantly, inflammatory activation, has been shown to promote TNTs between dendritic cells^38^, mesenchymal stem cells and cardiomyocytes^39^, and monocytes^40^. In addition, we and others have previously demonstrated that cells burdened with aggregate-prone proteins, including the exposure of neuronal and microglial cells to α-Syn aggregates, promotes the formation of intercellular connections^26,29,41–43^, although the underlying mechanism has remained elusive.

In this study, using human neuronal and microglial cell lines as well as iPSC-derived dopaminergic neurons (hNeurons) and microglia (hMG), we provide direct evidence that mitochondrial damage – induced by α-Syn aggregates – triggers STING-dependent innate immune activation, which in turn promotes the formation of TNTs and facilitates functional intercellular communication between neuronal and microglial cells. We identify a gate-keeping functionality of the major inflammatory transcription factor NF-κB in enabling the transfer of α-Syn from neuronal cells to microglia. Importantly, we demonstrate that damaged mitochondria are also transferred from α-Syn-burdened neuronal cells to microglia, where they are targeted for degradation, revealing an unrecognized dimension of microglial neuroprotection. While previous studies have described mitochondrial donation from microglia to neurons as a protective mechanism^29,32^, our data uncover the reciprocal transfer of damaged organelles from neurons to glia as a complementary response to cellular stress. This work provides the first mechanistic evidence that neurodegeneration-associated inflammation actively drives TNT-mediated neuron–microglia communication, highlighting a previously uncharacterized role for this pathway in modulating early neuroimmune interactions in disease contexts.

## Results

### 𝜶-Syn aggregates localize to mitochondria and disrupt their morphology

We previously reported that exogenous α-Syn aggregates largely localize with lysosomes in human neuronal cell line SH-SY5Y, and human microglial cell line HMC3^44^. Given α-Syn has affinity for mitochondrial membranes, we first examined whether exogenously added aggregates also associate with mitochondria. Following 16h of aggregate exposure, we observed that approximately 10% of mitochondria (immunostained for TOMM20) overlapped with α-Syn in both SH-SY5Y and HMC3 cells (**Fig. 1a, b**). A similar phenotype was seen in human iPSC-derived hNeurons and hMG, where ∼20% of mitochondria overlapped with aggregates (**Fig. 1c, d**). Structured-illumination super-resolution microscopy and 3D reconstruction confirmed these associations as direct contacts between aggregates and the mitochondrial outer membrane (**Fig. 1e, Extended Data** Fig. 1a). We also observed a time-dependent increase in mitochondrial association with α-Syn when cells were exposed to aggregates for increasing durations ranging from 1h until 12h (**Extended Data** Fig. 1b, c, d). Since α-Syn aggregates are internalized and trafficked with lysosomes during this time course^44^, we investigated how they reach mitochondria. Previous studies have shown that α- Syn disrupts lysosomal integrity, causing lysosome membrane permeabilization (LMP)^27,44^. We hypothesized that lysosomal escape enables aggregates to interact with mitochondria. Supporting this, treatment with the LMP-inducing agent L-leucyl-L-leucine methyl ester (LLOMe)^45^ led to a significant increase in α-Syn–TOMM20 overlap (**Extended Data** Fig. 1e**– g**), consistent with lysosomal rupture facilitating mitochondrial association (**Extended Data** Fig. 1h).

**Fig. 1.**
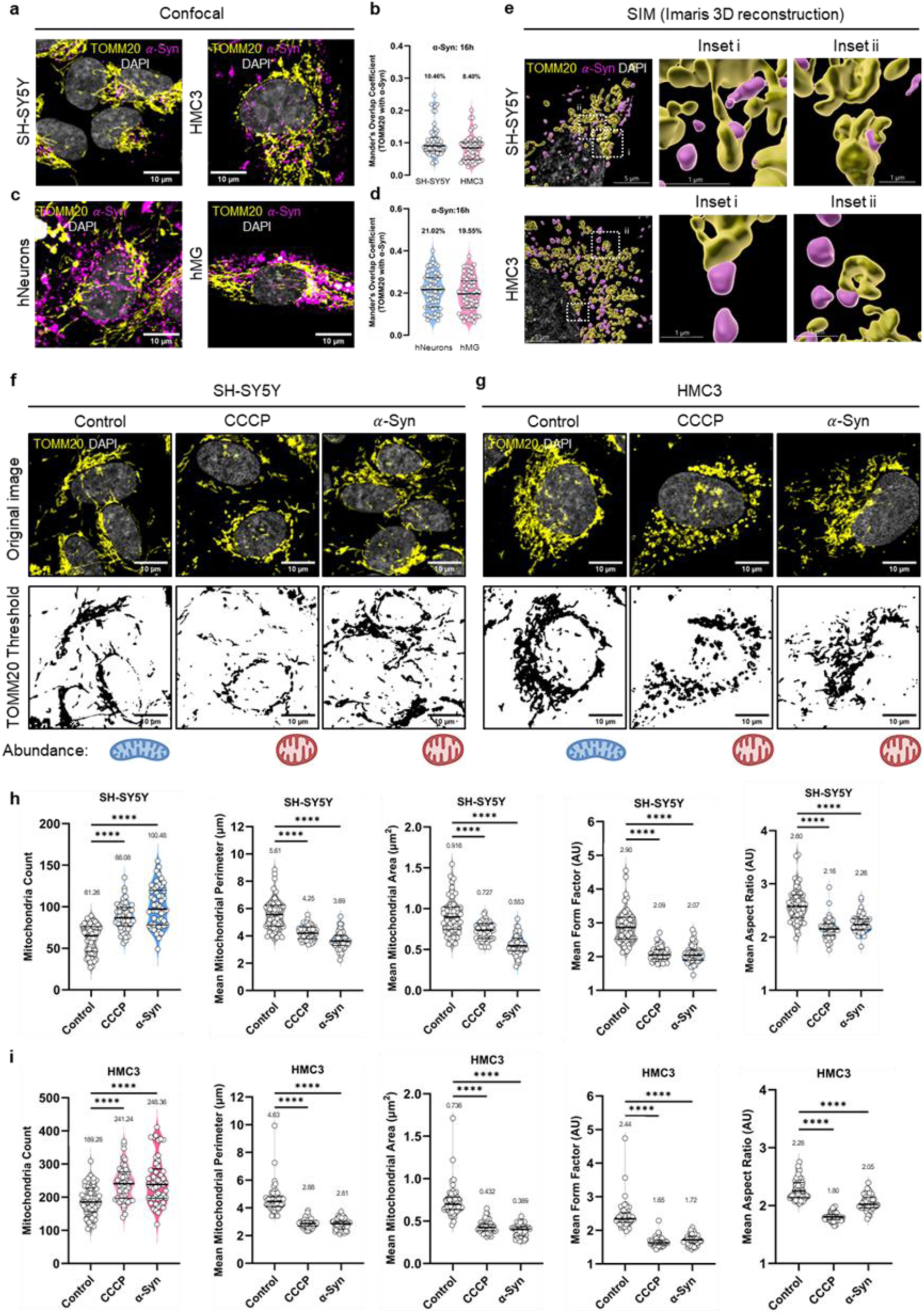
𝜶**-Syn localization to mitochondria and morphological alterations.** (a) Representative confocal images of α-Syn overlap with TOMM20+ mitochondria in neuronal cells (left panels) and microglial cells (right panels). (b) Quantification of Mander’s overlap coefficient of fraction overlap between mitochondria and α-Syn. N=3 independent experiments, n=50 cells. (c) Representative confocal images of α-Syn overlap with TOMM20+mitochondria in hiPSC-derived neurons (hNeurons; left panels) and microglia (hMG; right panels). (d) Quantification of Mander’s overlap coefficient of fraction overlap between mitochondria and α-Syn. N=3 independent experiments, n=60 cells. (e) Super-resolution images depicting overlap of α-Syn and mitochondria in neuronal cells (top panels) and microglial cells (bottom panels). (f-g) Morphological analysis of TOMM20 positive mitochondria using plug-in thresholded images (bottom panels) generated from original raw images (top panels) for neuronal cells (f) and microglial cells (g). Relative abundance of major observable mitochondrial phenotype depicted by the cartoon below. (h-i) Morphological quantifications of mitochondrial numbers (count), perimeter, area, form factor, and aspect ratio per cell for neuronal cells (h) and microglial cells (i). N=3 independent experiments, n=50 cells. Statistical significance was calculated using Brown-Forsythe and Welch ANOVA tests with Dunnett’s T3 multiple comparison. ****p<0.0001. Data represented as median and quartiles, with mean values mentioned within the graphs.

α-Syn has been reported to alter the morphology of mitochondrial network in several different manners^46^. To assess whether α-Syn-mitochondria association in our system lead to similar structural changes, we performed morphometric assessment of TOMM20-stained mitochondria (**Fig. 1f, g**). Compared to control cells, treatment with the oxidative phosphorylation uncoupler CCCP or exposure to α-Syn for 16h resulted in an increase in the number of TOMM20+ mitochondrial particles per cell with reduced perimeter and area. In addition, the form factor and aspect ratio of these particles were also reduced, indicative of mitochondrial fragmentation (**Fig. 1h, i and Extended Data** Fig. 1i).

### **𝜶**-Syn causes mitochondrial dysfunction

To determine whether these morphological alterations were associated with mitochondrial dysfunction, we next assessed mitochondrial membrane potential (ΔΨm), which is essential for maintaining oxidative phosphorylation and ATP production. A previous study on primary rat neuron-astrocyte co-culture reported depolarization of mitochondrial membrane upon exposure to α-Syn oligomers^47^. To test this in our model we used tetramethylrhodamine methyl ester (TMRM) as a readout of ΔΨm in SH-SY5Y and HMC3 cells exposed to α-Syn or CCCP (**Fig. 2a**). Expectedly, CCCP treatment significantly reduced ΔΨm by over 75% in both SH- SY5Y and HMC3 cells (**Fig. 2b, c**). Notably, the impact of α-Syn was much robust in neuronal cells (near 70% depolarization relative to control) than in microglia (∼29% depolarization), indicative of heightened vulnerability of neuronal mitochondria towards α-Syn aggregates-induced dysfunction.

**Fig. 2.**
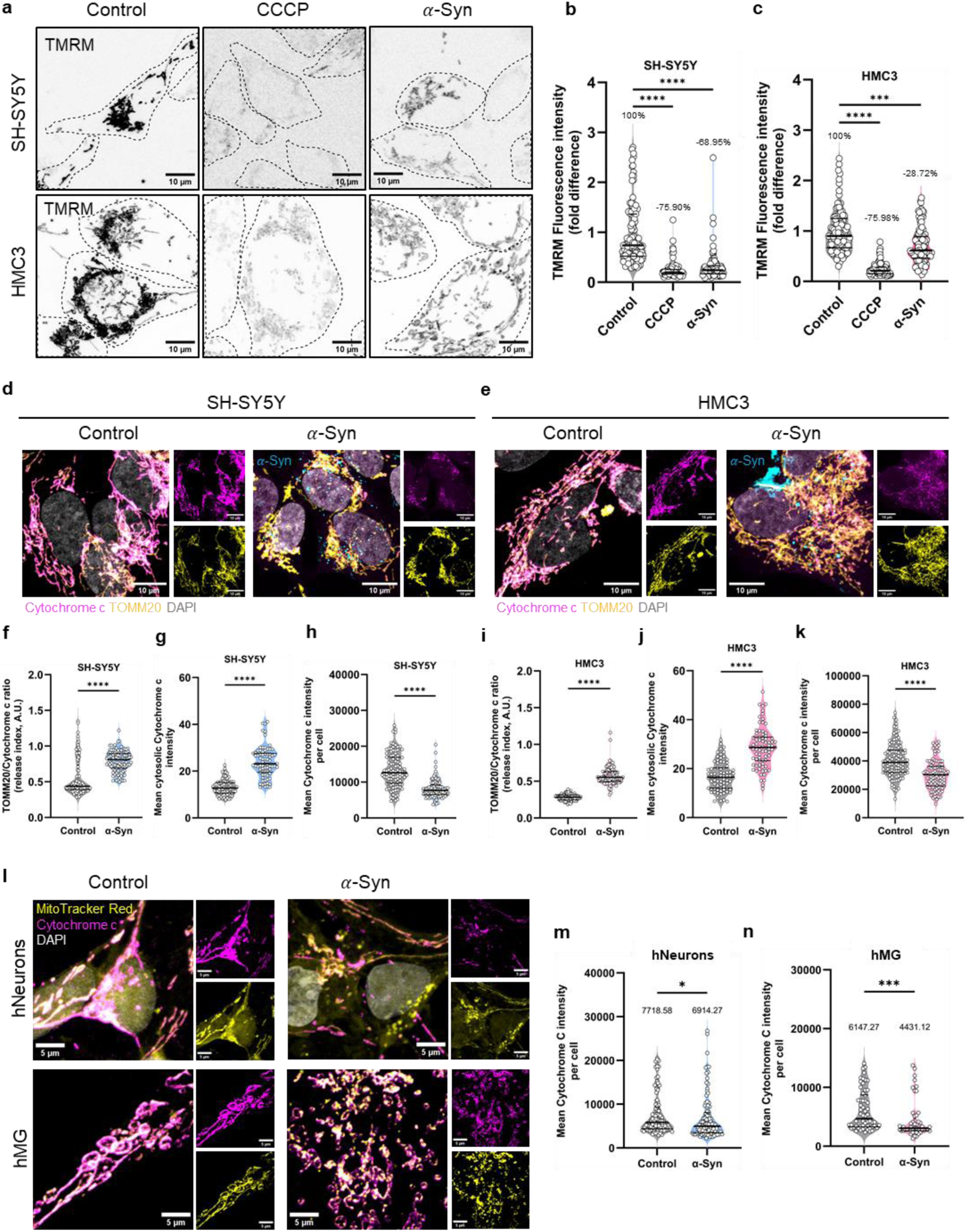
𝜶**-Syn exposure compromises mitochondrial functionality.** (a) Representative images of TMRM-stained neuronal cells (top panels) and microglial cells (bottom panels). (b- c) Quantification of fold difference of mean fluorescence intensities of TMRM in neuronal cells (b) and microglial cells (c). N=3 biological experiments, n=100 cells. Statistical significance was analyzed using Kruskal-Wallis test with Dunn’s multiple comparison. ***p<0.001, ****p<0.0001. (d-e) Representative confocal images of Cytochrome c level in neuronal cells (d) and microglial cells (e). (f) Cytochrome c release index (mean TOMM20 intensity/mean Cytochrome c intensity) in neuronal cells. N=3 independent experiments, n=100 cells. (g) Mean cytosolic Cytochrome c intensity in neuronal cells. N=3 independent experiments, n=75 cells. (h) Mean Cytochrome c intensity per cell in neuronal cells. N=3 independent experiments, n=100 cells. (i) Cytochrome c release index (mean TOMM20 intensity/mean Cytochrome c intensity) in microglial cells. N=3 independent experiments, n=100 cells. (j) Mean cytosolic Cytochrome c intensity in microglial cells. N=3 independent experiments, n=75 cells. (k) Mean Cytochrome c intensity per cell in microglial cells. N=3 independent experiments, n=100 cells. For (f-k), statistical significance was analyzed using Mann-Whitney test. ****p<0.0001. (l) Representative confocal images of Cytochrome c level in iPSC-derived neurons (hNeurons) and microglia (hMG). (m-n) Quantification of mean Cytochrome c intensity per cell for hNeurons (m) and hMG (n). N=3 independent experiments, n=100 cells for hNeurons, n=58 control and 49 α-Syn-treated hMG cells. Statistical significance was analyzed using Mann-Whitney test. *p<0.05, ***p<0.001. Data in all the graphs are represented as median and quartiles.

Mitochondrial depolarization has been demonstrated to be associated with release of Cytochrome c, an intermembrane space protein^48^, although depolarization can occur significantly later than Cytochrome c release into the cytoplasm^49^. To determine whether the observed loss in neuronal and microglial ΔΨm was also accompanied by Cytochrome c release, we performed co-immunostaining with TOMM20 (**Fig. 2d, e**). We observed significant Cytochrome c release in both cell types upon exposure to α-Syn aggregates (**Fig. 2f, i**), accompanied by elevated cytosolic levels (**Fig. 2g, j**). and a reduction in overall Cytochrome c protein levels (**Fig. 2h, k**). Taken together, these results are indicative of outer membrane permeabilization (MOMP) and functional impairment of mitochondria following α-Syn exposure. Similar experiments in hNeurons and hMG also revealed a reduction of Cytochrome c levels (**Fig. 2l, m, n**), underscoring the detrimental impact of α-Syn on mitochondrial functionality in both cell lines and iPSC-derived cells.

Having observed significant mitochondrial damage, we next asked whether aggregate- exposed cells were capable of clearing damaged mitochondria via mitophagy, a process implicated to be impaired in several neurodegenerative diseases^50^. Dysfunctional mitochondria are targeted to autophagosomes via interactions of mitophagy receptors with LC3 (mitophagosomes), which then fuse with lysosomes (mitolysosomes) for degradation. To address mitophagy flux in neuronal and microglial cells following α-Syn exposure, we performed co-immunostaining against TOMM20 and LC3 (to detect mitophagosomes; **Extended Data** Fig. 2a), and against TOMM20 and LAMP1 (to detect mitolysosomes; **Extended Data** Fig. 2d) in the presence or absence of the autophagy flux inhibitor bafilomycin A1 (Baf A1). Upon α-Syn exposure we detected an increase in mitophagosomes in both cell types (**Extended Data** Fig. 2b, c). However, Baf A1 treatment further elevated mitophagosome levels only in microglial cells, and not in neuronal cells. A similar phenotype was observed for mitolysosomes (**Extended Data** Fig. 2e, f) suggesting impaired mitophagy in neuronal cells but preserved or elevated flux in microglia. This is in accordance with our previous observations of impaired autophagy flux in neuronal cells but not in microglia following α-Syn exposure^44^, consequentially affecting mitophagy dynamics.

### **𝜶**-Syn leads to mtDNA release

Permeabilization of the outer mitochondrial membrane (MOMP) enables the release of mitochondrial matrix components, including mtDNA into the cytosol. Having observed MOMP in both neuronal and microglial cells upon α-Syn exposure, we next asked whether this led to mtDNA release. Co-immunostaining for double-stranded DNA (dsDNA) (to label mtDNA) and mitochondrial transcription factor A (TFAM) revealed the presence of both TFAM-negative mtDNA and TFAM-positive nucleoids outside mitochondria (**Fig. 3a**). The proportion of extra- mitochondrial mtDNA increased significantly in both cell types upon α-Syn exposure (**Fig. 3b, c**). 3D reconstructions of mtDNA and mitochondrial fluorescence signals further confirmed this, enabling quantification of mtDNA retained within mitochondria (DNA-in) versus released in the cytosol (DNA-out) (**Fig. 3d, e**). Such events were accompanied by the assembly of activated BAX (6A7) on the mitochondrial membrane (**Fig. 3f, g, h**), consistent with reports that BAX macropores facilitate mtDNA efflux^51^.

**Fig. 3.**
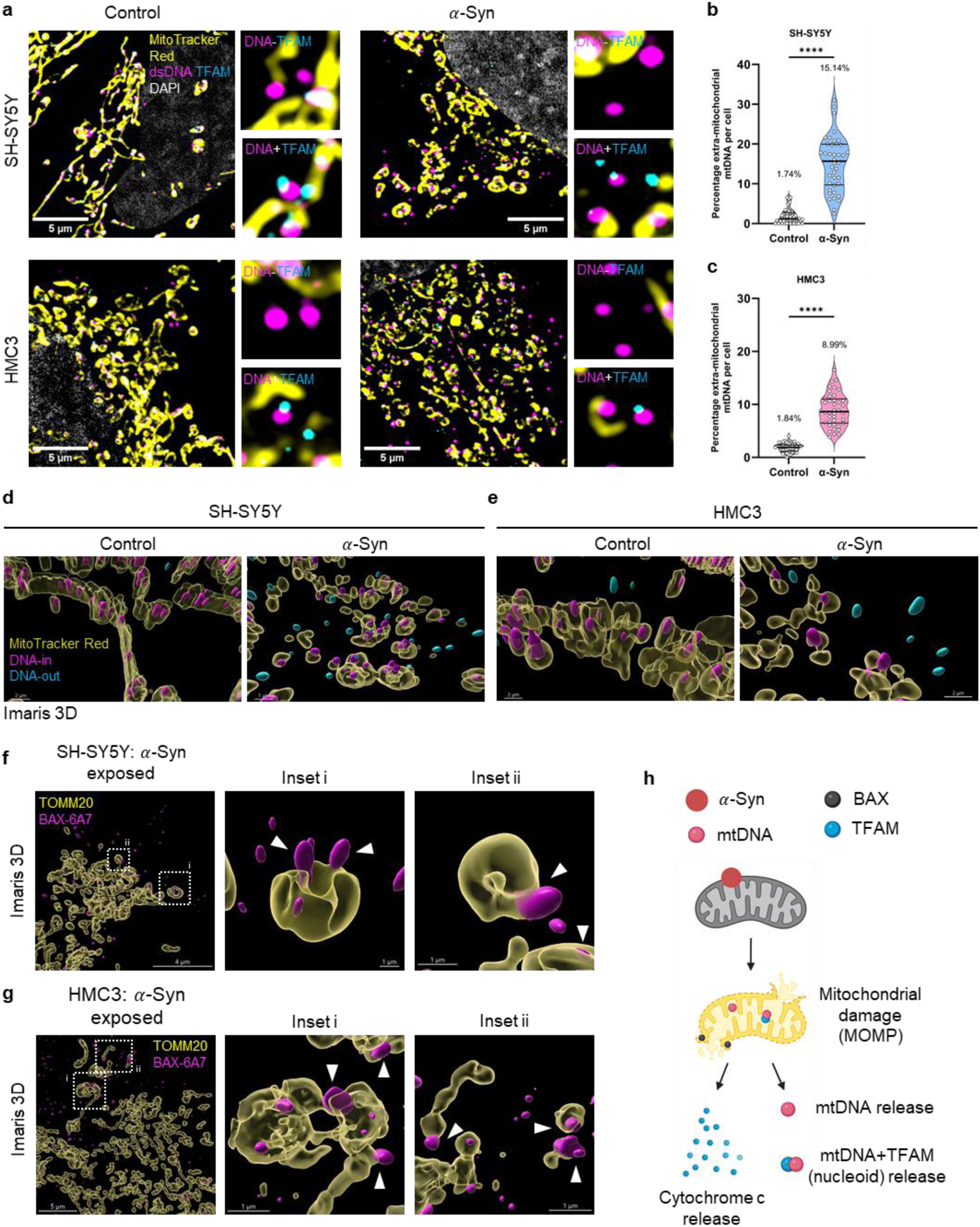
𝜶**-Syn induces mtDNA release in the cytoplasm.** (a) Representative super- resolution SIM images of neuronal cells (top panels) and microglial cells (bottom panels) labeled for mitochondria and stained for mtDNA and TFAM. (b-c) Quantification of extra- mitochondrial mtDNA per cell for neuronal cells (b) and microglial cells (c). N=3 independent experiments, n=40 cells. Statistical significance was analyzed using Mann-Whitney test. ****p<0.0001. Data represented as median and quartiles, with mean values mentioned within the graphs. (d-e) Imaris 3D reconstructions of depicting mtDNA encapsulated within mitochondria (magenta) or extra-mitochondrial (blue) for neuronal cells (d) and microglial cells (e). (f-g) Imaris 3D reconstruction of activated BAX (6A7) assembly on mitochondria in α-Syn-exposed neuronal cells (f) and microglial cells (g). (h) Schematic depicting the mechanism of mtDNA release in α-Syn-burdened cells.

### **𝜶**-Syn activates STING pathway and inflammation

Because of its ancestral prokaryotic origin, mtDNA release into the cytosol can trigger innate immune responses and consequent neuroinflammation. Upon recognition by cytosolic nucleic acid sensors such as cGAS, the second messenger cGAMP is produced, promoting dimerization and Golgi translocation of the ER-resident protein STING, marking its activation^52^. Exposure of both neuronal and microglial cells to α-Syn led to a marked increase in STING fluorescence intensity on GM130+ Golgi (**Fig. 4a, b, c**). A similar STING-Golgi overlap was observed in iPSC-derived hNeurons and hMG, suggesting a conserved biological response to α-Syn aggregates in different model systems (**Fig. 4d, e, f**). Following STING activation, Tank Binding Kinase 1 (TBK1) is recruited to the Golgi compartments and activated via transphosphorylation^53,54^. Consistently, we observed an increase in the number of pTBK1 puncta per cell (**Fig. 4g, h, i**), and the total area of these particles (**Fig. 4j, k**), confirming activation of the STING-TBK1 axis.

**Fig. 4.**
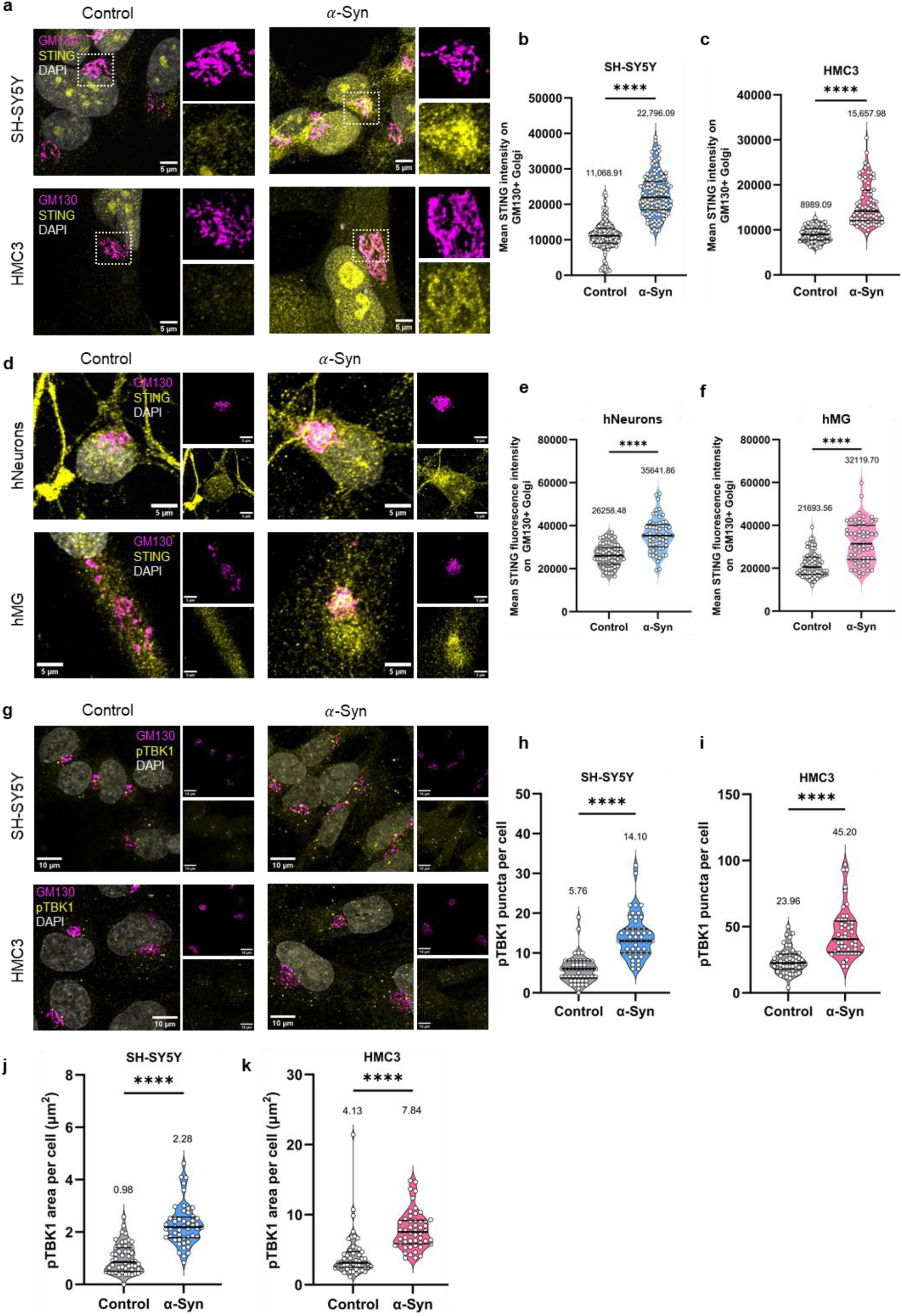
STING activation in. 𝜶**-Syn-exposed cells.** (a) Representative confocal images of STING translocation to GM130+ Golgi in neuronal cells (upper panels) and microglial cells (lower panels). (b-c) Quantification of mean STING fluorescence intensity on GM130+ Golgi in neuronal cells (b) and microglial cells (c). N=3 independent experiments; for neuronal cells: n=140 cells for control and 141 cells for α-Syn group, and for microglial cells: n=85 cells for control and 82 cells for α-Syn group. Statistical significance was analyzed using Mann-Whitney test. ****p<0.0001. (d) Representative confocal images of STING translocation to GM130+ Golgi in hiPSC-derived neurons (hNeurons; upper panels) and microglia (hMG; lower panels). (e-f) Quantification of mean STING fluorescence intensity on GM130+ Golgi in hNeurons (e) and hMG (f). N=3 independent experiments; n=60 cells per group. Statistical significance was analyzed using Mann-Whitney test. ****p<0.0001. (g) Representative confocal images of pTBK1 in neuronal cells (upper panels) and microglial cells (lower panels). (h-i) Quantification of the number of pTBK1 puncta per cell in neuronal cells (h) and microglial cells (i). (j-k) Quantification of pTBK1 area per cell in neuronal cells (j) and microglial cells (k). For (h-k), N=3 independent experiments, n=50 cells per group. Statistical significance was analyzed using Mann-Whitney test. ****p<0.0001. For all the graphs, data are represented as median and quartiles, with mean values mentioned within the graphs.

We next examined whether activation of the STING pathway led to an innate immune response, and whether this response differed between neuronal and microglial cells. Immunostaining for the key pro-inflammatory transcription factors downstream of STING – NF-κB and IRF3 – revealed a marked increase in nuclear localization of both total and phosphorylated NF-κB in response to α-Syn aggregates (**Extended Data** Fig. 3a–f). This activation was observed in both SH-SY5Y and HMC3 cell lines, as well as in iPSC-derived hNeurons and hMG (**Extended Data** Fig. 3g, h), indicating a conserved response across model systems. Similarly, total and phosphorylated IRF3 showed significantly enhanced nuclear accumulation in both cell types (**Extended Data** Fig. 3i–n). However, when we assessed downstream gene expression by RT-PCR, microglial cells exhibited a substantially stronger transcriptional response. Expression of pro-inflammatory cytokines (IL-1𝛽, TNF-α, IL-6, CD68; **Extended Data** Fig. 3o, p) and type I interferons and chemokines (IFNα, IFN𝛽, CXCL10, CCL20; **Extended Data** Fig. 3q, r) was consistently and significantly higher in HMC3 compared to SH-SY5Y cells. These results suggest that although both neuronal and microglial cells activate the STING pathway in response to α-Syn, microglia mount a much more robust innate immune response, highlighting their distinct roles in neuroinflammation and disease progression.

### **𝜶**-Syn-induced intercellular connections are dependent on inflammation

Based on our previous observation of α-Syn aggregates promoting intercellular connections in both neuronal and microglial cell lines^29^, and considering that neuroinflammation is a hallmark of neurodegenerative diseases including PD^55^, we hypothesized that inflammation may play a causative role in driving the formation of these connections. Since we observed significant release of mtDNA in both cell types in the presence of α-Syn aggregates (**Fig. 3**), we first asked whether cytosolic nucleic acid release alone could trigger intercellular connections. To test this, we treated SH-SY5Y and HMC3 cells with the BCL-2 family inhibitor ABT-737^56^, which induces limited MOMP and mtDNA release^57,58^ in combination with the pan-caspase inhibitor Q-VD-OPh. Upon treatment, both neuronal and microglial cells had significantly increased intercellular connections at all the time points of treatment (6, 12, and 24 hours) (**Extended Data** Fig. 4a, b, c), supporting the idea that mtDNA release is a key trigger for connection formation.

To then determine whether α-Syn-induced intercellular connections were dependent on innate immune activation, we inhibited key steps of the pathway. Since cytosolic mtDNA is sensed by cGAS, we first treated cells with the human cGAS-inhibitor G140^59^ alone or in combination with α-Syn aggregates (**Extended Data** Fig. 4d). As expected, α-Syn significantly increased the percentage of connected cells, but this effect was markedly reduced in the presence of G140 (**Extended Data** Fig. 4e, f), indicating a positive role of cGAS activity in the formation of intercellular connections. We next performed similar experiments using the STING inhibitor H-151 (prevents palmitoylation and clustering of STING)^60^ (**Fig. 5a**), and NF-κB inhibitor JSH- 23 (prevents its nuclear translocation)^61^ (**Fig. 5d**). In both cases, inhibition of these effectors significantly reduced the α-Syn-induced increase in intercellular connections (**Fig. 5b, c**, **5e, f**). Taken together, these results demonstrate that the cGAS–STING–NF-κB axis plays a positive regulatory role in the formation of intercellular connections in response to α-Syn aggregates (**Fig. 5j – left panel** and **Extended Data** Fig. 4g).

**Fig. 5.**
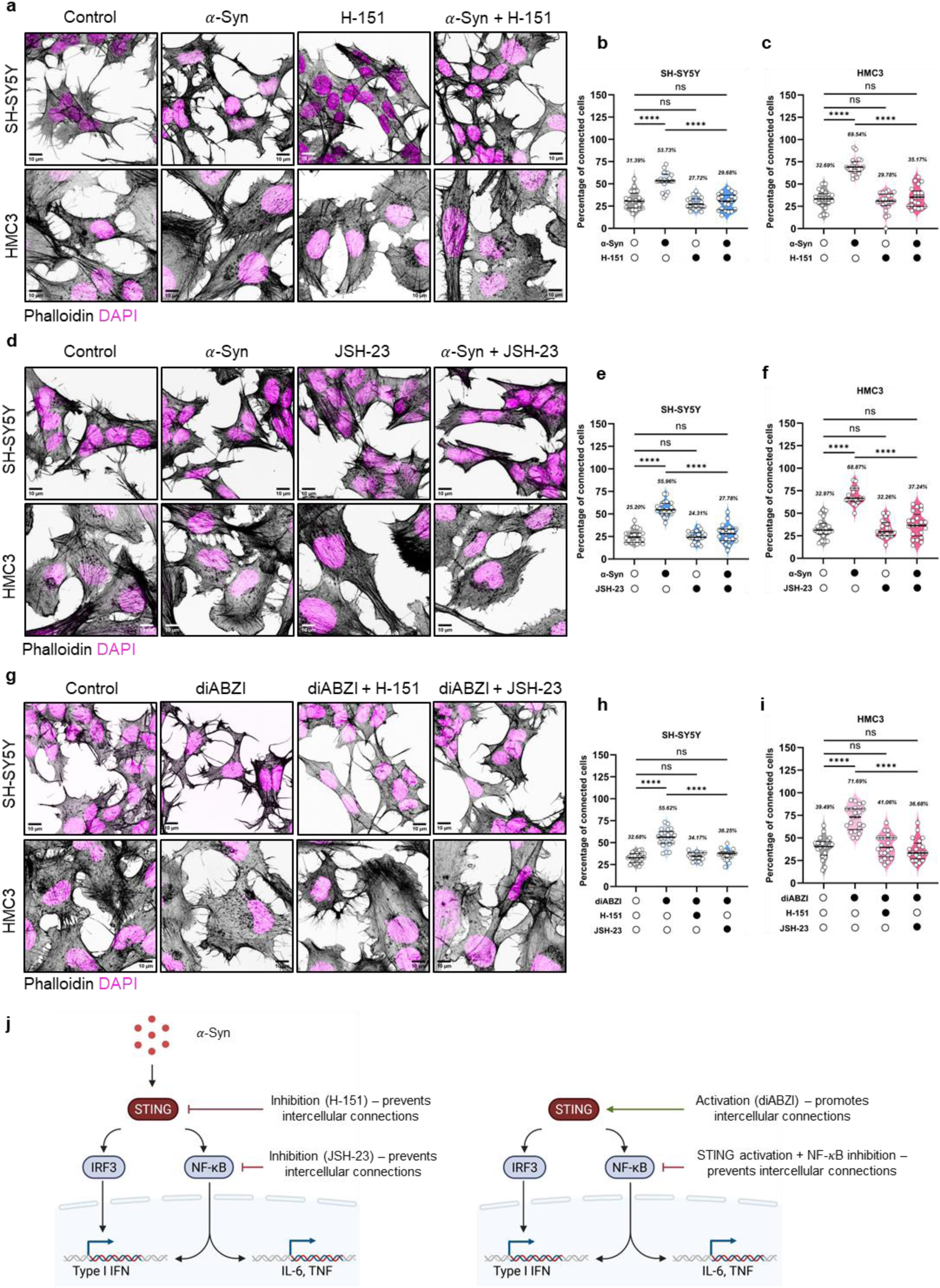
𝜶**-Syn-induced intercellular connections are inflammation-dependent.** (a) Representative phalloidin-stained images of neuronal cells (top panels) and microglial cells (bottom panels) treated with α-Syn aggregates, the STING inhibitor H-151, or both to assess for intercellular connections. (b-c) Quantification of the percentage of connected cells in different conditions for neuronal cells (b) and microglial cells (c). N=3 independent experiments, n=20-21 fields of views. Statistical significance was analyzed using Brown-Forsythe and Welch One-Way ANOVA and Dunnett’s T3 multiple comparison. ns: p>0.05, ****p<0.0001. (d) Representative phalloidin-stained images of neuronal cells (top panels) and microglial cells (bottom panels) treated with α-Syn aggregates, the NF-κB inhibitor JSH-23, or both to assess for intercellular connections. (e-f) Quantification of the percentage of connected cells in different conditions for neuronal cells (e) and microglial cells (f). N=3 independent experiments, n=18-21 fields of views. Statistical significance was analyzed using Brown- Forsythe and Welch One-Way ANOVA and Dunnett’s T3 multiple comparison. ns: p>0.05, ****p<0.0001. (g) Representative phalloidin-stained images of neuronal cells (top panels) and microglial cells (bottom panels) treated with the STING activator diABZI alone, or alongside H-151 or JSH-23 to assess for intercellular connections. (h-i) Quantification of the percentage of connected cells in different conditions for neuronal cells (h) and microglial cells (i). N=3 independent experiments, n=20-22 fields of views. Statistical significance was analyzed using Brown-Forsythe and Welch One-Way ANOVA and Dunnett’s T3 multiple comparison. ns: p>0.05, ****p<0.0001. Data in all the graphs are represented as median and quartiles, with mean percentage mentioned within the graphs. (j) Schematic of α-Syn-induced inflammatory regulators of intercellular connections (left panel), and effect of STING activation on such connections, depicting a gate-keeping activity of NF-κB in the process.

To further validate the involvement of STING in promoting intercellular connections, we treated both neuronal and microglial cells with diABZI, a potent non-nucleic acid STING agonist^62^ (**Fig. 5g**). Treatment with diABZI significantly increased the number of intercellular connections in both cell types, an effect abolished by co-treatment with the STING inhibitor H-151. Notably, the diABZI-induced increase in connections was also prevented by the NF-κB inhibitor JSH-23 (**Fig. 5h, i**). These results support a direct role for STING activation in promoting intercellular connectivity, and identify NF-κB as a critical downstream effector with a gatekeeping role in this process (**Fig. 5j, right panel**).

### Inflammation arises intracellularly and is independent of cell-surface receptors

α-Syn aggregates released by neurons have been reported to bind and activate microglial cell surface Toll-like receptors (TLRs) TLR2^63^ and TLR4^64^. To assess whether the inflammatory responses upon α-Syn exposure, resulting intercellular connections, observed in our systems depended on these TLRs activation, we treated cells with α-Syn aggregates alone, or in the presence of TLR4 inhibitor TAK-242 and TLR2 inhibitor TL2-C29. While α-Syn exposure promoted intercellular connectivity, neither inhibitors reduced this effect (**Extended Data** Fig. 5a–c). We therefore hypothesized that inflammation in our system may instead result from intracellular activation of the STING pathway. To test this, we measured nuclear NF-κB levels following co-treatment with α-Syn and either STING or TLR inhibitors (**Extended Data** Fig. 5d). Only STING inhibition significantly reduced nuclear NF-κB accumulation (**Extended Data** Fig. 5e, f), indicating that at 16 hours post α-Syn exposure, the observed inflammation and enhanced connectivity are driven by STING activation rather than TLR signaling.

### Inflammation induced connections enable functional interactions between neuronal cells and microglia

A key criterion for classifying intercellular connections as TNTs is their ability to transfer materials between connected cells. We have previously demonstrated that α-Syn aggregates can transfer from neuronal cells to microglia via TNTs^29^. To assess whether STING activation and resulting inflammation lead to functional TNT connections between these two cell types, we co-cultured α-Syn-exposed SH-SY5Y neuronal cells (donors) with HMC3 microglia (acceptors) in the presence of STING activity modulators (**Fig. 6a**). We observed a significant increase in the extent of transfer (percentage of acceptor microglia positive for α-Syn) upon STING activation with diABZI, but not when STING was activated in the presence of its inhibitor H-151 (**Fig. 6b**). In addition, H-151 also reduced the number of transferred α-Syn aggregates (**Fig. 6c**). Such transfer happened majorly in a contact-dependent manner, as analysis of conditioned media demonstrated minimal contribution of secretion on α-Syn transfer (**Fig. 6d**).

**Fig. 6.**
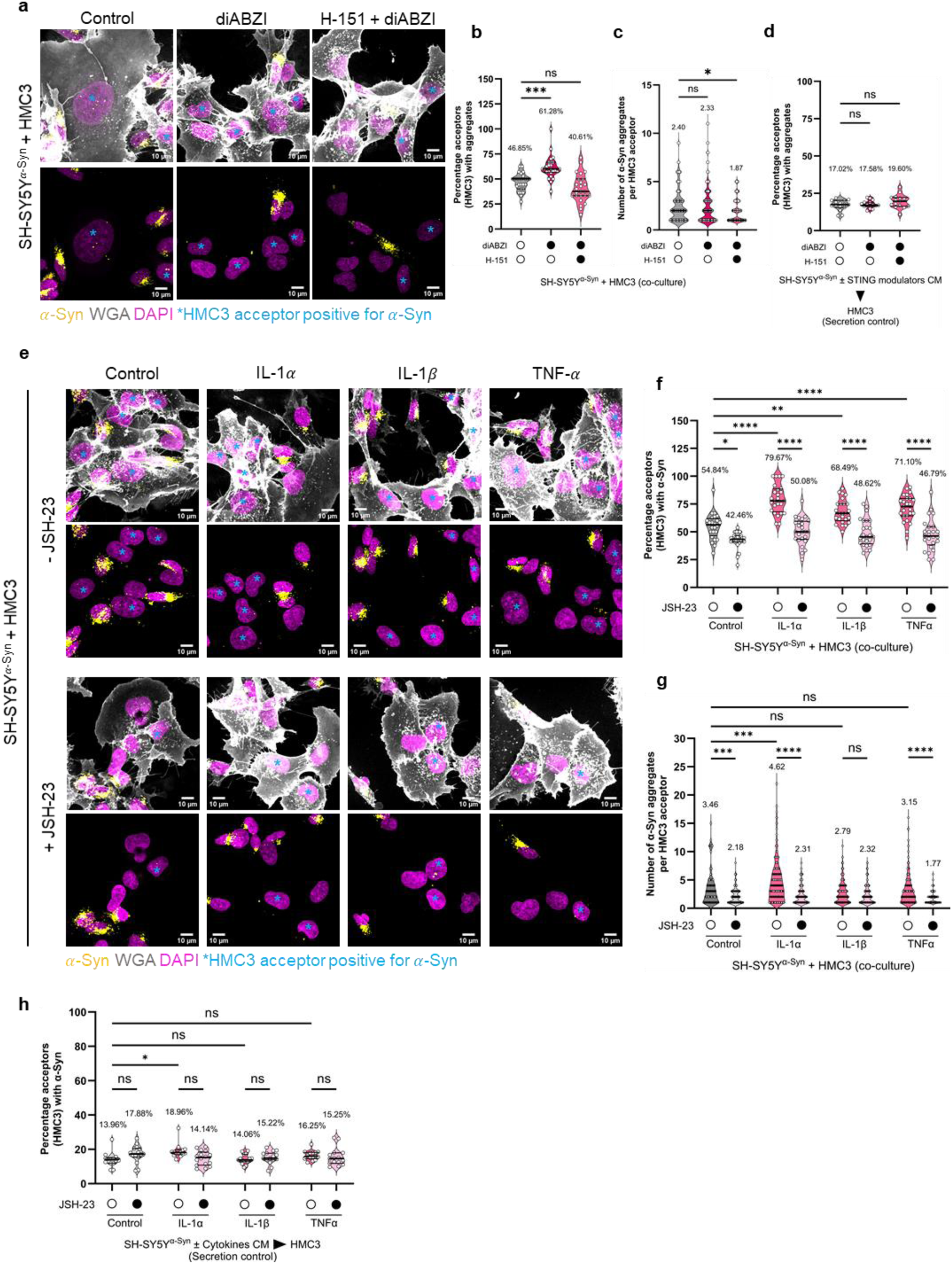
Inflammation promotes. 𝜶**-Syn spread to microglia via TNTs.** (a) Representative confocal images of neuronal cells loaded with α-Syn (donors) in co-culture for 24h with microglial cells (acceptors) in the presence of the STING activator diABZI alone, or alongside the STING inhibitor H-151. Cyan asterisks highlight acceptor microglial cells positive for α- Syn. (b) Quantification of the percentage of acceptor microglial cells positive for α-Syn. N=3 independent experiments, n=25-26 fields of views. Statistical significance was analyzed using Kruskal Wallis test with Dunn’s multiple comparison. ns: p>0.05, ***p<0.001. (c) Quantification of the number of α-Syn aggregates per acceptor microglial cell as in (b). n=106 cells for control, 177 cells for diABZI-treated group, and 92 cells for diABZI+H-151 co-treated group. Statistical significance was analyzed using Kruskal Wallis test with Dunn’s multiple comparison. ns: p>0.05, *p<0.05. (d) Quantification of the extent of α-Syn transfer via secretion (CM: conditioned media). N=3 independent experiments, n=15 fields of views. Statistical significance was analyzed using Kruskal Wallis test with Dunn’s multiple comparison. ns: p>0.05. (e) Representative confocal images of neuronal cells loaded with α- Syn (donors) in co-culture for 24h with microglial cells (acceptors) in the presence of pro-inflammatory cytokines alone (upper panels) or co-treated with the NF-κB inhibitor JSH-23 (lower panels). Cyan asterisks highlight acceptor microglial cells positive for α-Syn. (f) Quantification of the percentage of acceptor microglial cells positive for α-Syn. N=3 independent experiments, n=24-25 fields of views. Statistical significance was analyzed using Two-Way ANOVA with Šídák’s multiple comparison. *p<0.05, **p<0.01, ****p<0.0001. (g) Quantification of the number of α-Syn aggregates per acceptor microglial cell as in (f). n=126 (Control –JSH-23), 124 (control +JSH-23), 227 (IL-1α –JSH-23), 163 (IL-1α +JSH-23), 223 (IL-1𝛽 –JSH-23), 144 (IL-1𝛽 +JSH-23), 188 (TNFα –JSH-23), and 134 (TNFα +JSH-23) cells. Statistical significance was analyzed using Two-Way ANOVA with Šídák’s multiple comparison. ns: p>0.05, ***p<0.001, ****p<0.0001. (h) Quantification of the extent of α-Syn transfer via secretion (CM: conditioned media). N=3 independent experiments, n=14-16 fields of views. Statistical significance was analyzed using Two-Way ANOVA with Šídák’s multiple comparison. ns: p>0.05, *p<0.05. Data in all the graphs are represented as median and quartiles. Mean values are mentioned within the graphs.

Building on our observation of upregulation of cytokine expression in α-Syn-exposed cells (**Extended Data** Fig. 3o-r), we further explored the role of NF-κB in TNT formation. Treatment of neuronal and microglial mono-cultures with pro-inflammatory cytokines IL-1α, IL-1𝛽 and TNF-α increased the percentage of connected cells for both cell types. However, this cytokine-induced increase was abolished in the presence of the NF-κB inhibitor JSH-23 (**Extended**

**Data Fig. 6a, b, c, d**). To assess the functional nature of these connections, we performed co-culture experiments as above, where we assessed the extent of α-Syn transfer from neuronal cells (donors) to microglia (acceptors) in the presence of inflammatory modulators (**Fig. 6e**). While NF-κB activation through the aforementioned pro-inflammatory cytokines promoted α-Syn transfer to microglial cells, its inhibition significantly reduced the extent (**Fig. 6f**) and the number of transferred aggregates (**Fig. 6g**). Since α-Syn secretion remained minimal and largely unaffected by NF-κB modulation, our results support a direct, positive role for NF-κB in promoting TNT-mediated α-Syn transfer from neuronal cells to microglia.

TNT formation is linked to actin remodeling^65^. To understand how inflammation affects actin organization in promoting functional intercellular connections, we examined total and phosphorylated cofilin levels in cells treated with α-Syn, TNF-α, or cotreated with JSH-23. Actin depolymerizing factor (ADF)/cofilin promotes actin depolymerization and filament severing^66^, while its phosphorylation inactivates this function, favoring longer actin filaments in cells. Upon inflammatory stimulation (α-Syn and TNF-α), both neuronal and microglial cells showed increased total cofilin (**Extended Data** Fig. 6e, g), and an even stronger increase in phosphorylated cofilin (**Extended Data** Fig. 6f, h). Interestingly, this effect was abolished upon co-treatment with JSH-23. Taken together, these results suggest that inflammatory conditions promote F-actin stabilization through cofilin inactivation, supporting the formation of TNT-like structures.

To understand whether molecular changes also associated with global re-organization of the actin cytoskeletal network, we measured the coherency of actin filaments^67^ in experimental conditions where α-Syn-induced intercellular connections were shown to depend on NF-κB (**Fig. 5d**). A higher coherency of actin filaments would be expected upon directional re- organization of the network (**Extended Data** Fig. 6i), a phenotype expected during the formation of TNTs. We observed an increased actin coherency upon α-Syn treatment, which was lost upon NF-κB inhibition (**Extended Data** Fig. 6j, k), consistent with the reduced intercellular connections seen in **Fig 5e, f**. Taken together, these results indicate that inflammatory stress induces global actin remodeling that supports TNT formation and facilitates α-Syn transfer from neurons to microglia.

### Neuronal cells transfer damaged mitochondria to microglia

While a recent study elegantly demonstrated a neuroprotective function of microglia via TNT-mediated transfer of functional mitochondria^32^, it remains unclear whether neurons also transfer mitochondria to microglia. Given our findings of extensive mitochondria damage in α- Syn-treated neuronal cells (**Fig. 2**), we hypothesized that they are capable of transferring damaged mitochondria to microglia. To test this, we co-cultured naïve microglia and mitochondria-labeled neuronal cell (matrix-targeted DsRed – MitoDsRed) loaded or not with α-Syn (**Fig. 7a**). We observed a more than two-fold increase in the extent of mitochondrial transfer from aggregate-burdened neuronal cells, compared to control cells, with most of the transfer occurring in a contact-dependent manner (**Fig. 7b**). Additionally, microglia received significantly more mitochondrial particles from α-Syn-stressed neuronal cells (**Fig. 7c**).

**Fig. 7.**
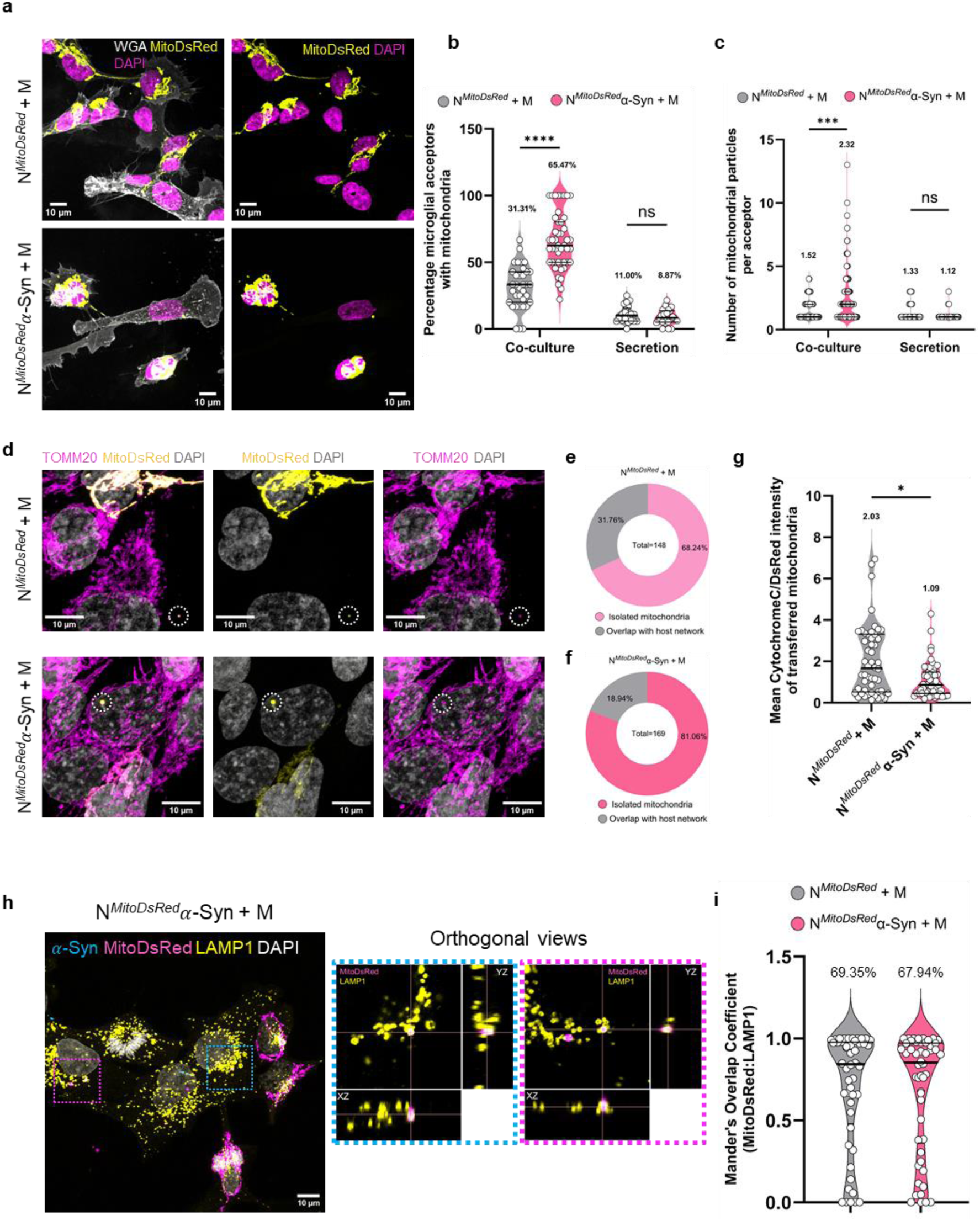
Neuronal cells transfer damaged mitochondria to microglial cells for degradation. (a) Representative confocal images of mitochondria-labeled control (upper panels) or α-Syn-loaded (bottom panels) neuronal cells (donors) in co-culture for 24h with microglial cells (acceptors). (b) Quantification of the percentage of microglial cells receiving MitoDsRed+ mitochondria in both co-cultures and conditioned media (secretion) experiments. N=3 independent experiments, n=35 (N*^MitoDsRed^* + M co-culture), 47 (N*^MitoDsRed^*α-Syn + M co- culture), 21 (N*^MitoDsRed^* + M secretion) and 22 (N*^MitoDsRed^*α-Syn + M secretion) fields of views. Statistical significance was analyzed using Two-Way ANOVA with Uncorrected Fisher’s LSD multiple comparison. ns: p>0.05, ****p<0.0001. (c) Quantification of the number of mitochondrial particles received per acceptor microglia as in (b). N=74 (N*^MitoDsRed^* + M co- culture), 185 (N*^MitoDsRed^*α-Syn + M co-culture), 30 (N*^MitoDsRed^* + M secretion), and 34 (N*^MitoDsRed^*α- Syn + M secretion) cells. Statistical significance was analyzed using Two-Way ANOVA with Uncorrected Fisher’s LSD multiple comparison. ns: p>0.05, ***p<0.001. (d) Representative confocal images of neuronal cell-derived MitoDsRed+ mitochondria in microglial cells. (e-f) Quantification of the proportion of transferred mitochondria from naïve neuronal cells (e) or α- Syn-burdened neuronal cells (f) that remain isolated in microglial cells, or overlap with the host network. Total number of mitochondrial particles analyzed is mentioned within the pie-chart. (g) Quantification of the relative health of transferred mitochondria (Cytochrome c level, indicative of outer membrane permeabilization state of mitochondria) from naïve (N*^MitoDsRed^* + M) or α-Syn-burdened (N*^MitoDsRed^*α-Syn + M) neuronal cells. N=3 independent experiments, n=50 transferred mitochondrial particles analyzed. Statistical significance was tested using Mann-Whitney test. *p<0.05. (h) Representative confocal image, and orthogonal views of dashed boxes, highlighting encapsulation of MitoDsRed+ mitochondrial particles derived from α-Syn-burdened neuronal cells within LAMP1+ lysosomes of microglial cells. (i) Quantification of overlap between MitoDsRed+ mitochondria and LAMP1+ lysosomes in acceptor microglial cells. Data in violin plots are represented as median and quartiles, with mean values mentioned within the graphs.

While functional mitochondria transfer from microglia to neuronal cells exert a neuroprotective effect via integration into the host mitochondrial network^32^, most mitochondria transferred from neuronal cells to microglia remained isolated and did not integrate in the microglial mitochondria network (**Fig. 7d, e, f**). The proportion of isolated mitochondria also increases from ∼68% when coming from naïve neuronal cells to ∼81% when derived from α-Syn- burdened neuronal cells. Measurements of Cytochrome c levels in these transferred organelles revealed that mitochondria derived from aggregate-burdened neuronal cells had a lower level of the protein (**Fig. 7g**), suggesting that outer membrane-permeabilized mitochondria (damaged) are preferentially transferred to microglial cells.

Given that the transferred mitochondria are damaged and do not integrate to the host mitochondrial network, we hypothesized that these organelles are destined for degradation, representing an alternative neuroprotective mechanism by which microglia clear neuronal-derived damaged components. Upon immunostaining for lysosomes, we observed significant engulfment of MitoDsRed particles within LAMP1+ vesicles in microglial cells, with the majority of mitochondrial particles targeted for clearance (**Fig. 7h, i**; median>0.84 for both groups). Taken together, these results demonstrate that microglia not only receive damaged mitochondria from α-Syn–stressed neuronal cells but also actively clear them, adding a new layer to microglial neuroprotective functions.

### **𝜶**-Syn-burdened neuronal cells prime microglia towards inflammation

Having observed mitochondrial damage-induced mtDNA release in α-Syn-burdened cells, and elevated mitochondrial transfer from stressed neuronal cells to microglia, we next asked whether the transferred mitochondrial particles contained mtDNA. To address this, we co-cultured SH-SY5Y-MitoDsRed cells (donors) with acceptor HMC3 cells, and immunostained for both TOMM20 and DNA (**Fig. 8a**). We found that majority of transferred mitochondrial particles were mtDNA+ (63.51% in control group, versus 75.81% upon α-Syn exposure) (**Fig. 8b**). Notably, we also observed mtDNA extruding from the mitochondrial matrix after reaching microglial cells (**Fig. 8a – inset iii**). As such, we next assessed the inflammatory state of microglial cells when co-cultured with naïve or α-Syn-burdened neuronal cells. We observed a mild activation of STING in microglial cells co-cultured with aggregate-laden neuronal cells (**Fig. 8c, d**), along with increased nuclear levels of NF-κB (**Fig. 8e, f**) and IRF3 (**Fig. 8g, h**) in such cells. Interestingly, this inflammatory response was independent of the number of α-Syn aggregates received by microglia (**Extended Data** Fig. 7a, b, c). However, microglial cells had reduced Cytochrome c levels when co-cultured with α-Syn-burdened neuronal cells (**Extended Data** Fig. 7d). Taken together, these results suggest a bystander priming of microglia towards inflammation, through either mtDNA release in microglial cells, alterations of microglial mitochondrial functionalities, and/or other secretory factors.

**Fig. 8.**
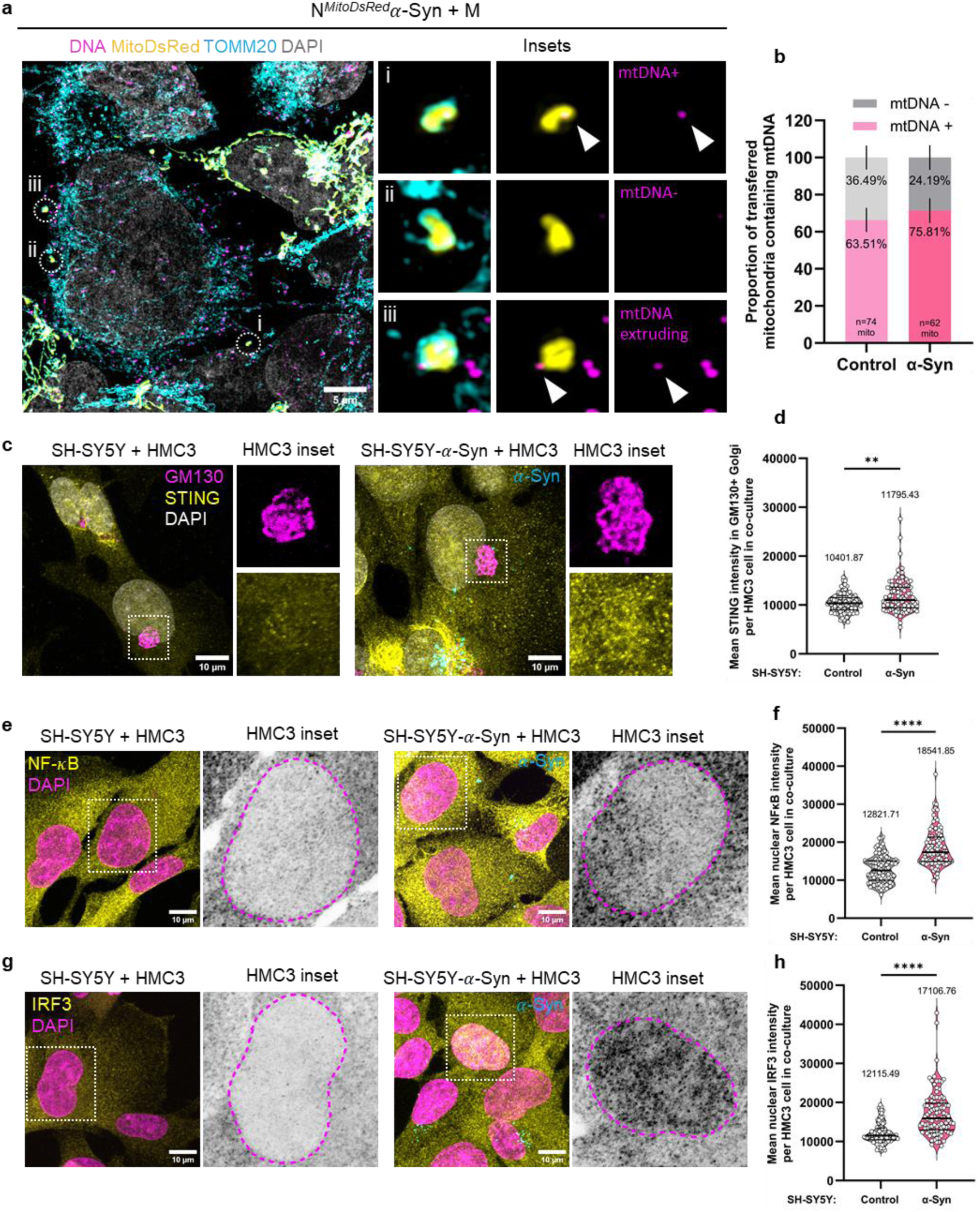
mtDNA status of transferred mitochondria and bystander inflammation. (a) Representative super-resolution SIM images of α-Syn-burdened MitoDsRed+ neuronal cells in co-culture with microglial cells for 24h. Insets i, ii, and iii highlight transferred mitochondrial particles that are mtDNA+, mtDNA-, and have mtDNA extruding (white arrowhead) from the matrix, respectively. (b) Quantification of the proportion of transferred mitochondria from control or α-Syn-burdened neuronal cells that are positive or negative for mtDNA. N=3 independent experiments, numbers of mitochondrial particles analyzed are mentioned within the bars. Error bars represent SEM. (c) Representative confocal images of STING translocation to GM130+ Golgi in microglial cells when co-cultured for 24h with naïve (left panels) or α-Syn-burdened (right panels) neuronal cells. (d) Quantification of mean STING fluorescence intensity on GM130+ Golgi in microglial cells. N=3 independent experiments, n=100 cells. Statistical significance was analyzed using Mann-Whitney test. **p<0.01. (e) Representative confocal images of nuclear translocation of NF-κB in microglial cells when co- cultured for 24h with naïve (left panels) or α-Syn-burdened (right panels) neuronal cells. (f) Quantification of mean nuclear NF-κB fluorescence intensity in microglial cells. N=3 independent experiments, n=100 cells. Statistical significance was analyzed using Mann- Whitney test. ****p<0.0001. (g) Representative confocal images of nuclear translocation of IRF3 in microglial cells when co-cultured for 24h with naïve (left panels) or α-Syn-burdened (right panels) neuronal cells. (h) Quantification of mean nuclear IRF3 fluorescence intensity in microglial cells. N=3 independent experiments, n=100 cells. Statistical significance was analyzed using Mann-Whitney test. ****p<0.0001. Data in violin graphs are represented as median and quartiles. Mean values are mentioned within the graphs.

## Discussion

The roles of TNTs in mediating intercellular communication between neuronal and glial cells have been very well established. Neuronal cells can transfer α-Syn aggregates to astrocytes and microglia via TNTs, enabling their clearance^28,44^. Moreover, microglia have been shown to exert neuroprotective effects through mitochondrial transfer to neurons^29,32^. However, a central unsolved question in the field pertains to the molecular signals that drive the formation of TNTs under Synucleinopathic stress. Here, we uncover a previously unrecognized role of innate immune activation—specifically the cGAS–STING–NF-κB signaling axis—in driving TNT-mediated intercellular communication between neuronal cells and microglia in response to α-Syn aggregates (**Extended Data** Fig. 7e).

Exogenous α-Syn aggregates are efficiently internalized by both neuronal and microglial cells, and accumulate predominantly in lysosomes in both murine and human neuronal cells^26,27,44^, as well as in human microglia^44^, leading to extensive lysosomal damage. Besides lysosomes, α-Syn has been reported to have affinities for other sub-cellular membranes, such as that of mitochondria in both cell cultures and dopaminergic neurons of SNpc^68^. Oligomeric complexes of α-Syn have been demonstrated to bind to TOMM20 and disrupt protein import into mitochondria^15^. α-Syn aggregates also promote neuronal death via inhibition of ATP production and mitochondrial permeability transition pore (mPTP) opening^47^. In addition, localization of α-Syn with mitochondria externalizes cardiolipin, a major anionic lipid of the inner mitochondrial membrane, which in turn promotes further seeding of α-Syn aggregates^69,70^. All these results hint towards a major biological role of mitochondria in mediating α-Syn pathology.

In our study, using exogenous pre-formed fibrils of α-Syn, we demonstrate that these aggregates can escape from damaged lysosomes and associate with mitochondria. This association leads to a significant compromise in mitochondrial integrity – manifested by organelle fragmentation – and function – as evidenced by loss of membrane potential and release of Cytochrome c. Notably, these effects were more pronounced in neuronal cells than in microglia, suggesting a greater vulnerability of neuronal mitochondria. The persistence of damaged mitochondria in neuronal cells likely reflects impaired clearance due to reduced mitophagy flux, consistent with our previous observation of compromised autophagy in these cells^44^. In contrast, microglial cells may compensate through enhanced mitochondrial biogenesis—though this remains to be tested—enabled by the elevated mitophagy flux observed, thereby maintaining mitochondrial homeostasis more effectively.

As a consequence of mitochondrial damage and outer membrane permeabilization, we observed significant release of mtDNA in the cytoplasm, a phenomenon previously reported in several neurodegenerative diseases, including Parkinson’s^71^, Alzheimer’s^21,72^, and amyotrophic lateral sclerosis^25^. In our system, ectopic mtDNA triggered a robust inflammatory response in both neuronal and microglial cells by activating the STING pathway which in turn induced the expression of pro-inflammatory cytokines and type I interferons. This innate immune activation ultimately promoted the formation of intercellular connections driven by the global re-organization of the actin cytoskeleton. Although previous studies have demonstrated an interplay between inflammatory triggers and TNTs^38–40^, the molecular mechanism driving such long TNT connections under these conditions have remained largely undefined. Importantly, we identify this inflammatory activation as the driver of TNT formation through NF-κB–dependent actin remodeling. We also report increased levels of phospho-Cofilin, which impairs actin depolymerization, promoting the formation of long actin filaments essential for TNT-like structures. Although the upstream regulators of cofilin phosphorylation remain to be identified, our findings define inflammation as a mechanistic trigger for TNT biogenesis under Synucleinopathic stress.

In line with the downstream consequences of mtDNA release, we identify the STING pathway as a positive regulator of functional TNTs between neuronal cells and microglia. Notably, NF-κB—a central pro-inflammatory transcription factor downstream of STING—appears to act as a gatekeeper, ultimately determining whether intercellular connections are formed. We further show that the spread of α-Syn aggregates from neuronal cells to microglia is significantly enhanced in pro-inflammatory environments triggered by STING activation or cytokine treatments. Importantly, this enhanced transfer can be reversed by pharmacological inhibition of STING (H-151) or NF-κB (JSH-23), underscoring the critical role of inflammation in promoting α-Syn propagation. These findings are consistent with previous reports implicating inflammatory signaling in facilitating α-Syn spread *in vivo*^73,74^. The implications of cGAS-STING pathway activation in α-Syn pathology spread *in vivo*, however, remains to be determined.

In addition to aggregate transfer, we observe that damaged mitochondria are also transferred from α-Syn–burdened neuronal cells to microglia, where they are subsequently degraded. Given the extensive mitochondrial damage induced by α-Syn and the impaired mitophagy observed in neuronal cells, TNT-mediated transfer to microglia likely serves as a compensatory mechanism to maintain cellular homeostasis. In our earlier work, we reported increased autophagic flux in bystander microglia co-cultured with α-Syn–stressed neurons^44^. Here, we also demonstrate bystander inflammation in these microglia, which may further promote autophagy induction through NF-κB–mediated signaling^75,76^. Such an inflammatory response in microglial cells could be arising because of a pro-inflammatory environment caused by α-Syn-burdened neuronal cells, mtDNA extrusion from transferred mitochondria, instigated mtDNA release from microglial mitochondria, or via gap junctions-mediated transfer of cGAMP, which has previously been reported to spread bystander inflammation^77,78^.

Collectively, our findings suggest that neuronal reliance on microglia for the clearance of damaged mitochondria represents a biologically rational mechanism underlying mitochondrial damage-induced TNT formation. This study reveals, for the first time, a moonlighting role for mitochondria—not only as targets of dysfunction but also as active initiators of innate immune signaling that promotes intercellular communication and the propagation of pathogenic α-Syn aggregates. This finding adds a critical new layer to our understanding of neurodegenerative disease progression and opens new avenues for targeting TNT-mediated pathology spread.

## Materials and Methods

### Culture of Cell lines

Human neuroblastoma cell line SH-SY5Y (referred to as neuronal cells in the manuscript) were cultured in RPMI1640 media (Euroclone, ECB2000L), and the human microglia clone 3 cell line HMC3 (referred to as microglial cells) were cultured in DMEM (Sigma-Aldrich, D6429), both supplemented with 10% foetal calf serum (FCS) (Eurobio Scientific, CVFSVF00-01) and 1% penicillin-streptomycin (Gibco, 15140-122; 100 units/mL final concentration). Cells were cultured in a humidified CO2 incubator at 37°C, and passaged using 0.05% Trypsin-EDTA (Gibco, 25300-054). Cells were counted before each experiment, and seeded on UV-treated 12 mm glass coverslips (uncoated) (Epredia, CB00120RA120MNZ0) for fixed-cell imaging, and on 35 mm glass-bottom microdishes (Ibidi GmbH, 81156) for live-cell imaging. Cells between passages 3 and 10 were used for all the experiments.

### Culture of hiPSC lines

All procedures adhered to Spanish and EU guidelines and regulations for research involving the use of human pluripotent cell lines. The human iPSC lines used in our studies were generated following procedures approved by the Commission on Guarantees concerning the Donation and Use of Human Tissues and Cells of the Carlos III Health Institute, Madrid, Spain.

Control hiPSC line SP11 was used for neurons, and SP13wt/wt for microglia. Generation and characterization of these lines have been described elsewhere^79,80^. hiPSC were maintained on Matrigel (Corning, 354234)-coated plastic plates (Thermo Fisher Scientific) and in mTeSRTM1 medium (Stem Cell Technology, 85850) until the start of the protocols of differentiation.

### Microglia generation

The employed protocol for the differentiation of microglia and its characterization is as previously described^81^. Briefly, hiPSC were passed in single colonies and after 2-4 days, mTeSRTM1 medium was supplemented with 80 ng/mL of Bone Morphogenetic Protein (BMP)-4 (PeproTech, 120-05) for a total of 4 days. From the following day, cells were changed using SP34 medium (StemProTM-34 SFM (Gibco™, 10639011), 1% of P/S (Cultek, SV30010) and 1% of Ultraglutamine (Glut; Lonza, LZBE17-605EU1). For two days, media was supplemented with 80 ng/ml of VEGF (PeproTech, AF-100-20), 100 ng/ml of Stem Cell Factor (SCF, PeproTech, 300-07) and 25 ng/ml of Fibroblast Growth Factor (FGF)-2 (PeproTech, 100-18B). From days 7 to 14, supplemented factors included 50 ng/ml of Fms-like tyrosine kinase 3-Ligand (Flt3-L, Humanzyme, HZ-1151), 50 ng/ml of IL-3 (PeproTech, 200-03), 50 ng/ml of SCF, 5 ng/ml of Trombopoietin (TPO, PeproTech, 300-18) and 50 ng/ml of M-CSF (PeproTech, 300-25). The last step consisted on the addition to SP34 medium of Flt3-L, M-CSF and 25 ng/ml of GM-CSF (PeproTech, 300-03), with media changes every 3-4 days. Starting from day 35, floating microglial progenitors were collected from the culture’s supernatant and passed through a 70 μm Filcon™ Syringe-Type nylon mesh (BD Biosciences, 10271120). Cells were counted and centrifuged at 300xg for 10 minutes. Recollected progenitor microglial cells were plated at a final density of 5,000 microglia per well of a 24 well plate (Thermo Fisher Scientific) and on top of plastic coverslips. Media was changed twice a week with RPMI 1640 Medium (GibcoTM, 11875093) supplemented with 50 ng/mL of IL-34 (PeproTech,200-34) and M-CSF. Microglia were considered mature after one week in culture.

### Neuron generation

NPCs were generated following a previously published protocol from^82,83^. Briefly, iPSC was split into a 96 well-plate, V-bottom shape and centrifuged 800xg for 10 minutes to force their aggregation. Cells were grown on mTeSR medium (STEMCELL technologies, 05825) for 24 hours. Embryoid bodies (EBs) were plated in a 60mm dish and the medium was then changed to Proneural [DMEM/F12 (Life, 21331-20) and Neurobasal (Life, 21103-049) – 1:1, 0.5% N2 (Life, 17502048), 1% B27 w/o Vitamin A (Life, 12587-010), 1% L-Glutamine (Linus, X0551-100), 1% Penicillin/Streptomycin (ScienCell, 0503), and 2-Mercaptoethanol (gibco 31350- 010)]. EBs were seeded in POLAM-coated (poly-L-ornitine Sigma-aldrich, P4957; laminin, Sigma-Aldrich, L2020) wells of a 6 well plate with Proneural supplemented with Noggin 200ng/ml (PeproTech, 120-10C) and SB431542 10 µM (TOCRIS, 301836-41-9). When NEP rosettes were visible, they were enzymatically dissociated with trypsin 0.05% to obtain NPCs and plated on POLAM-coated wells of a 12 well plate. NPCs were then split up to 6-8 times in order to purify the culture.

DAn were differentiated following a previously published protocol^84^. NPCs at 80-100% confluency were cultured on POLAM-coated (poly-L-ornithine Sigma-aldrich, P4957; laminin, Sigma-Aldrich, L2020) wells in DAn induction medium [DMEM/F12 (Life, 21331-20), 1% N2 (Life, 17502048), 1% Penicillin/Streptomycin (ScienCell, 0503)] supplemented with 200 ng/ml Sonic Hedgehog (PeproTech, 100-45) and 100 ng/ml FGF8 (PeproTech, 100-25) for 6 days. This step allowed for NPCs patterning towards dopaminergic fate. DAn progenitors were then plated on POLAM-coated dishes in N2B27 medium (DMEM/F12 (Life, 21331-20) – Neurobasal (Life, 21103-049) 1:1, 0.5% N2 (Life, 17502048), 1% B27(Life, 17504-044), 1% L-Glutamine (Linus, X0551-100), and 1% Penicillin/Streptomycin (ScienCell, 0503) for 10 days for maturation. For terminal differentiation DAn were cultured on Matrigel (Corning, 354234) coated wells supplemented with 20 ng/ml BDNF (Peprotech, 450-02) and 20 ng/ml GDNF (Peprotech, 450-10) for 25 days.

### Preparation of ***𝜶***-Syn aggregates

α-Syn aggregates were prepared as described before^85^. Briefly, human wild-type α-Syn was purified from Escherichia coli BL21 (DE3) with RP-HPLC. Using a manufacturer’s protocol of labeling kit, fibrils were either conjugated with Alexa Fluor™ 488, 568, 647 fluorophores (Invitrogen), or not, and stocks were stored at -80°C for long-term, and -20°C for short-term storage and immediate use. Prior to exposure to cells, fibrils were diluted in growth medium at a working concentration of 500 nM and sonicated (BioBlock Scientific, Vibra Cell 7504) for 5 minutes at an amplitude of 80%, pulsed for 5 seconds “on” and 2 seconds “off”. For iPSC-derived cells, owing to their fragile nature, a lower concentration of fibrils (200 nM) was used. Sonicated fibrils were then added on cells directly without the addition of any intracellular delivery agents for designated time points.

### Reagents and primary antibodies

For inflammatory modulations, following reagents were used with associated treatment concentration and durations: STING activator – diABZI (SelleckChem – compound 3, S8796; 2.5 𝜇M for 2h towards the end of experiment), STING inhibitor – H-151 (Invivogen InvitroFit^TM^, inh-h151; 0.5 𝜇g/mL for 16h for TNT assessment experiments, and 24h for α-Syn transfer experiments), NF-κB inhibitor – JSH-23 (Sigma, J4455; 10 𝜇M for 16h for TNT assessment experiments, and 24h for α-Syn transfer experiments), human cGAS inhibitor – G140 (Invivogen InvitroFit^TM^, inh-g140; 10 𝜇M for 16h), Bcl-2 inhibitor – ABT-737 (Sigma, 197333; 10 𝜇M for 6h, 12h, or 24h) co-treated with pan-caspase inhibitor – Q-VD-OPh (Non-O-methylated, Sigma, 551476, 10 𝜇M), human IL-1α (Sigma, I2778; 3 ng/mL for 24h), human IL-1𝛽 (Sigma, H6291; 20 ng/mL for 24h), human TNF-α (Merck Millipore, 654205; 50 ng/mL for 24h), TLR4 inhibitor – TAK-242 (Tocris, 6587; 10 𝜇M for 16h), TLR2 inhibitor – TL2-C29 (Invivogen InvitroFit^TM^, inh-C29; 100 𝜇M for 16h), lysosomotropic agent LLOMe (Sigma, L7393; 1 mM for 1h towards the end of experiment), mitochondrial electron transport chain uncoupler CCCP (Sigma, C2759; 20 𝜇M for 2h).

The following dyes were used: MitoTracker Red CMX Ros (Invitrogen, M7512; 500 nM for 30 minutes at 37°C) and TMRM (Invitrogen, T668; 100 nM for 30 minutes at 37°C).

Primary antibodies against the following antigens were used: TOMM20 (Santa Cruz, sc- 17764; 1:400), Cytochrome c (BD Pharmigen, 556432, 1:400), dsDNA (Progen, clone AC-30-10, 690014; 1:100), TFAM (Invitrogen, PA5-29571; 1:400), GM130 (BD Biosciences, 610823; 1:100), STING (Invitrogen, PA5-23381; 1:100), phospho-Ser172-TBK1 (Cell Signaling Technology, D52C2; 1:50), NF-κB (Thermo Fischer, 51-0500; 1:100), phospho-Ser536-NF-κB (Invitrogen, Ma5-15160, T.849.2; 1:100), IRF3 (Cell Signaling Technology, D6I4C; 1:100), phospho-Ser386-IRF3 (Cell Signaling Technology, E7J8G; 1:100), activated BAX-6A7 (Santa Cruz, sc23959; 1:100; kind gift from Julien Prudent), LC3B (Cell Signaling Technology, D11, 1:500), LAMP1 (DSHB, H4A3, 1:100), Cofilin (Invitrogen, PA5-17372; 1:1000 for western blotting), phospho-Ser3-Cofilin (Invitrogen, PA5-17752; 1:1000 for western blotting), GAPDH (Sigma, G9545; 1:5000 for western blotting).

### Immunocytochemistry

Immunofluorescence on cell lines were performed using a standard protocol, as described previously^44^. In brief, cells were fixed with 4% paraformaldehyde (PFA [Electron Microscopy Sciences, 15710]) for 30 minutes at room temperature (RT) and incubated in 50 mM ammonium chloride (NH4Cl [Sigma Aldrich, A0171]) solution for 15 minutes at RT to quench the fixative. Following three washes, cells were permeabilized with Triton-X100 [Sigma Aldrich, 9002-93-1] (0.1% v/v solution in 1X DPBS) for 5 minutes at RT. Cells were then incubated in blocking solution (2% w/v in 1X DPBS) for 1h at RT. Cells were then incubated with primary antibodies overnight (for 16h) at 4°C. The next day, cells were incubated with respective secondary antibodies (all from Thermo fisher Scientific, dilution of 1:500 in blocking solution) for 1h at RT. Cells were then washed three times with 1X DPBS, counterstained for nuclei with DAPI (1:1000 in PBS [Sigma-Aldrich, D9542]) and mounted on glass slides with Aqua-Poly/Mount (Polysciences Inc., 18606-20). Slides were imaged at least a day after mounting of coverslips.

hiPSC were fixed with 4% PFA for 20 minutes followed by permeabilization with 0.1 % Triton- X100 (Sigma-Aldrich, 9036-19-5) solution. Cells were blocked with a solution composed of 0.3% Triton-X100 and 3% normal donkey serum (Millipore, 41105901) diluted in TBS for 2h at RT. Cells were then incubated with primary antibodies in afore-mentioned dilutions overnight at 4°C. The next day, cells were re-blocked for 1h at RT, followed by secondary antibody incubation for 2h at RT. Finally, cells were incubated with DAPI (Abcam, 228549), diluted to 1:5000 in TBS), and coverslips were mounted on glass slides with PVA-DABCO (Sigma Aldrich) mounting media.

### Mitophagy flux analysis

To assess mitophagy flux upon exposure of cells to α-Syn aggregates, cells were immunostained for TOMM20 and LC3 (mitophagosomes), or TOMM20 and LAMP1 (mitolysosomes). To inhibit mitophagy flux, a saturating concentration of 400nM of Bafilomycin A1 (Sigma, SML1661) was added for 4h before fixation and immunostaining.

### qRT-PCR

RNA was isolated using TRIZOL reagent (Invitrogen) following manufacturer’s protocol. 1 μg of RNA was used to synthesize cDNA using high capacity cDNA reverse transcription kit (4368814, Applied Biosystems). qPCR was carried out using diluted cDNA (20 ng per reaction) using iTaq universal SYBR Green supermix (1725124, Bio-Rad). Relative mRNA expression was calculated using the ΔΔCt method and each gene was normalized with Ct value of β- actin. Three technical replicate reaction mixes were performed for each gene and biological replicate. Primers against the following human genes were used:

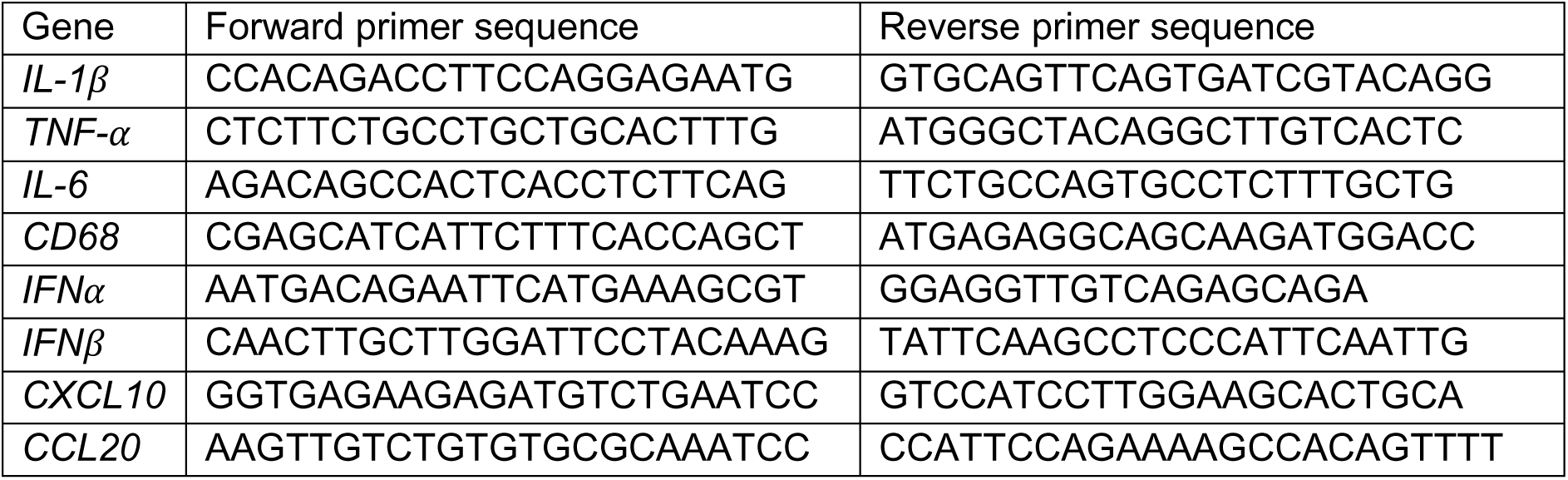

### Western blotting

Both SH-SY5Y and HMC3 cells were grown in 6-well dishes for 24h before appropriate treatment. Following incubation, cells were trypsinized and pellets were homogenized in 100 μl of radioimmunoprecipitation assay (RIPA) buffer consisting of 50 mM Tris-HCI (pH 7.4), 1% TritonX-100, 0.5% sodium-deoxycholate, 0.1% sodium dodecyl sulphate, 150 mM sodium chloride and 2 mM ethylenediaminetetraacetic acid, supplemented with 1X protease inhibitor (cOmplete Mini, EDTA-free; Sigma, 11836170001). Protein concentrations were measured using Bradford’s method, following manufacturer’s instructions (Thermo Fisher Scientific). Proteins (20 µg) were denatured in SDS (8%) and 𝛽-mercaptoethanol (5%) at 95 °C for 10 minutes. After separation by SDS-PAGE on a 3-8% tris-acetate gel, proteins were electro- transferred to nitrocellulose membranes. The membranes were blocked with 5% BSA and incubated overnight with primary antibodies (as per the aforementioned dilutions) in blocking buffer, followed by incubation with horseradish peroxidase-conjugated secondary antibody (Millipore, 1:5000) in TBS containing 0.1% Tween-20 at room temperature for 1h. Finally, proteins were visualized using an ECL kit (Thermo Fisher Scientific) and chemiluminescence images were acquired using GE Amersham Imager AI680 analyzer.

### ***𝜶***-Syn transfer assay

To assess for α-Syn transfer, neuronal cells or microglia were grown for 24h in a 6-well dish, and then neuronal cells were exposed to 500 nM of α-Syn. After 16h of incubation, both neuronal cells (now the donor population) and microglial cells (acceptor population) were trypsinized and seeded on 12 mm coverslips in a 1:1 ratio, and co-cultured for 24h. Inflammatory modulators were added for the entire co-culture duration, except for diABZI which was added for the final 2h. Cells were then fixed to preserve TNTs (described below) and stained with wheat germ agglutinin 647 (WGA, Thermo Fisher Scientific, W32466; 3.33 𝜇g/mL) for 15 minutes at RT, followed by nuclei counterstaining with DAPI (1:1000 in 1X DPBS).

### TNT counting

To efficiently visualize TNTs in cultures, cells were fixed at sub-confluency (∼70%) as per previous protocols^29,86^. In brief, two different fixative solutions were used – fixative 1 (0.05% glutaraldehyde [GA {Sigma Aldrich, G5882}], 2% PFA, 0.2 M HEPES buffer [Gibco, 15630- 080] in 1X DPBS), followed by fixative 2 (4% PFA, 0.2 M HEPES buffer in 1X DPBS) for 15 minutes each at RT. Cells were then labelled with Phalloidin 647 (Thermo Fisher Scientific, A12380; 1:250 in 1X DPBS) and DAPI (1:1000 in 1X DPBS) for 15 minutes each at RT, before mounting on glass slides.

### Fixed-cell microscopy

Images were acquired using Zeiss LSM900 inverted confocal microscope equipped with four lasers (wavelength in nm): 405, 488, 561 and 640 nm. For TNT counting, samples were imaged using 40X oil immersion objective (1.3 numerical aperture) with a field-of-view effective zoom of 0.8x, whereas all other immunofluorescence samples were acquired using a 63X oil immersion objective (1.4 numerical aperture) with a 1x zoom. Image acquisition was performed using ZEN blue software. The entire cell volume was imaged for all samples, with optical sections of 0.45 𝜇m. Depending on the cell types, the entire volume of cells ranged between 7 and 13 𝜇m in thickness.

Super-resolution images were acquired using Zeiss LSM 780 Elyra SIM set-up (Carl Zeiss, Germany) using Plan-Apochromat 63×/1.4 oil objective with a 1.518 refractive index oil (Carl Zeiss). 16-bit images were acquired in “frame-fast” mode between wavelengths, with appropriate grid sizes. Optical thickness was set at 0.133 𝜇m. Raw images were processed using the SIM processing tool of Zen black software.

### Quantification and Statistical Analyses

Colocalization analysis was performed using the JACoP plug-in FIJI^87^. For TNT counting, images were processed for analysis using the “manual TNT annotation” plug-in of ICY software (https://icy.bioimageanalysis.org/plugin/manual-tnt-annotation/). Mitochondrial morphology was analyzed using the mitochondria analyzer plug-in of FIJI^88^. Actin coherency was analyzed using OrientationJ plug-in of FIJI (developed by the Biomedical Imaging Group, EPFL, Switzerland). No prior power analysis was done to measure the sample size. Graphs were plotted using GraphPad prism 10.0, and appropriate statistical tests were performed on raw data. For all datasets, an initial normality distribution test was performed, and non-parametric tests were performed for any datasets that did not satisfy normal distribution.

Statistical tests performed are mentioned in the respective figure legends. All the experiments were performed for three independent biological replicates, unless stated otherwise.

## Supporting information

Supplementary Figures with legends

## Acknowledgments

We thank all the members of the Membrane Traffic and Pathogenesis Unit, Institut Pasteur, for insightful discussions. We thank Reine Bouyssie, a member of the administrative staff of the Membrane Traffic and Pathogenesis Unit for her continued support. We thank Prof. Julien Prudent for kindly providing BAX-6A7 antibody.

## Funding

Pasteur-Paris University International Doctoral Program (RC)

The Journal of Cell Science Travelling Fellowship JCSTF24101615 (RC) France Parkinson - Soutien de l’Association France Parkinson 2021 (CZ) Don Explore AD - Programme Explore de l’Institut Pasteur (CZ)

Agence Nationale de la Recherche ANR-20-CE13-0032-01 (CZ) Fondation pour la Recherche Médicale FRM - EQU202103012692 (CZ)

PID2022-139546OB-I00 supported by MCIN/AEI/10.13039/501100011033 and FEDER, and

PDC2021-121051-I00 supported by MCIN/AEI/10.13039/501100011033 and by the European Union Next Generation EU/ PRTR) (AC)

AGAUR (2021-SGR-974) (AC)

VT was the recipient of a pre-doctoral La Caixa INPhINIT Incoming Fellowship (code: LCF/BQ/DI21/11860038

JMM was the recipient of a pre-doctoral fellowship FPI (PRE2022-104573) from the Spanish Ministry of Economy and Competitiveness (MINECO).

Research was conducted within the context of Pasteur International Joint Research Unit Neurodegenerative Diseases.

## Author contributions

Conceptualization: RC, CZ; Methodology: RC, SM, VT, JMM, TN, MH; Investigation: RC, PS, VT, JMM; Visualization: RC; Supervision: CZ, AC; Funding acquisition: CZ, AC; Writing— original draft: RC; Writing—review & editing: RC, SM, VT, JMM, AC, CZ.

## Competing interests

Authors declare that there are no competing interests.

## Data and materials availability

All data are available in the main text, or in the supplementary materials.

## Notes

### Competing Interest Statement

The authors have declared no competing interest.

