## Supplementary Figures with legends for "α-Synuclein aggregates induce mitochondrial damage and trigger innate immunity to drive neuron–microglia communication"

### Extended Data Figures

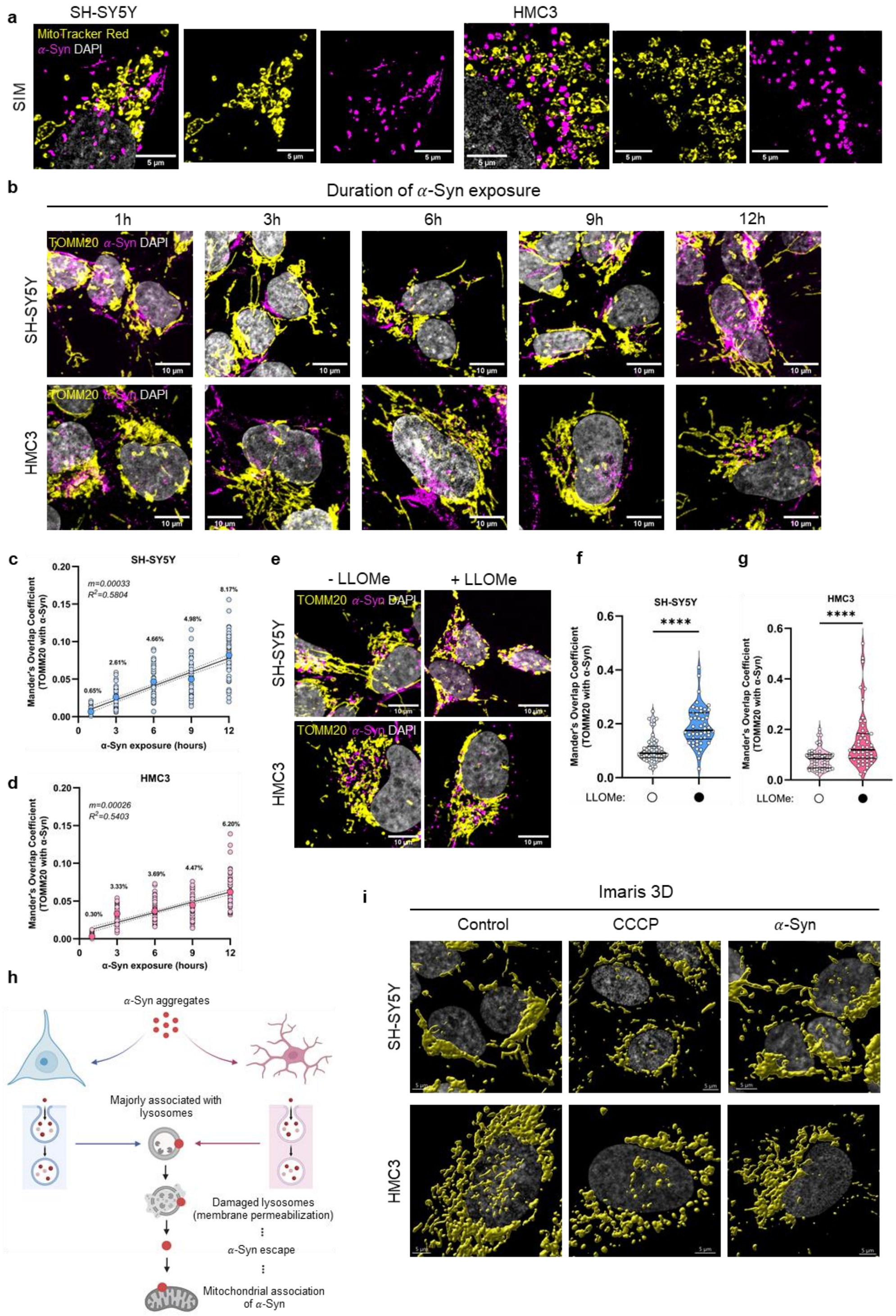


**Extended Data Fig. 1. $\alpha$-Syn localization with mitochondria, and morphological alterations (related to Main Fig. 1).** (a) Representative Imaris 3D reconstructions of super-resolution SIM images highlighting $\alpha$-Syn association with mitochondria in neuronal cells (left panels) and microglial cells (right panels). (b) Time-course analysis of $\alpha$-Syn association with mitochondria in neuronal cells (top panels) and microglial cells (bottom panels). (c-d) Quantification of Mander's overlap co-efficient of fraction of TOMM20+ mitochondria overlapping with $\alpha$-Syn per neuronal cell (c) and microglial cell (d). N=3 independent experiments, n=50 cells per time-point. Mean values for each time point are in bold, darker circles. Solid lines represent simple linear regression fit, with 95% confidence interval denoted by dashed lines. Mean percentage of overlap mentioned within the graphs. (e) Representative confocal images of neuronal cells (top panels) and microglial cells (bottom panels) treated or not with LLOMe to assess $\alpha$-Syn localization with TOMM20+ mitochondria. (f-g) Quantification of Mander's overlap coefficient of fraction of TOMM20+ mitochondria overlapping with $\alpha$-Syn after 16 hours of incubation in control and LLOMe-treated (for 1h) conditions in neuronal cells (f) and microglial cells (g). Control groups are same as in main Fig. 1b. N=3 independent experiments, n=50 cells. Statistical significance was analyzed using Mann-Whitney test. ****p<0.0001. Data represented as median and quartiles. (h) Schematic representation of lysosomal escape of $\alpha$-Syn and subsequent mitochondrial association. (i) Imaris 3D reconstruction of TOMM20+ mitochondria in control, CCCP-treated, and $\alpha$-Syn-exposed conditions for neuronal cells (top panels) and microglial cells (bottom panels).

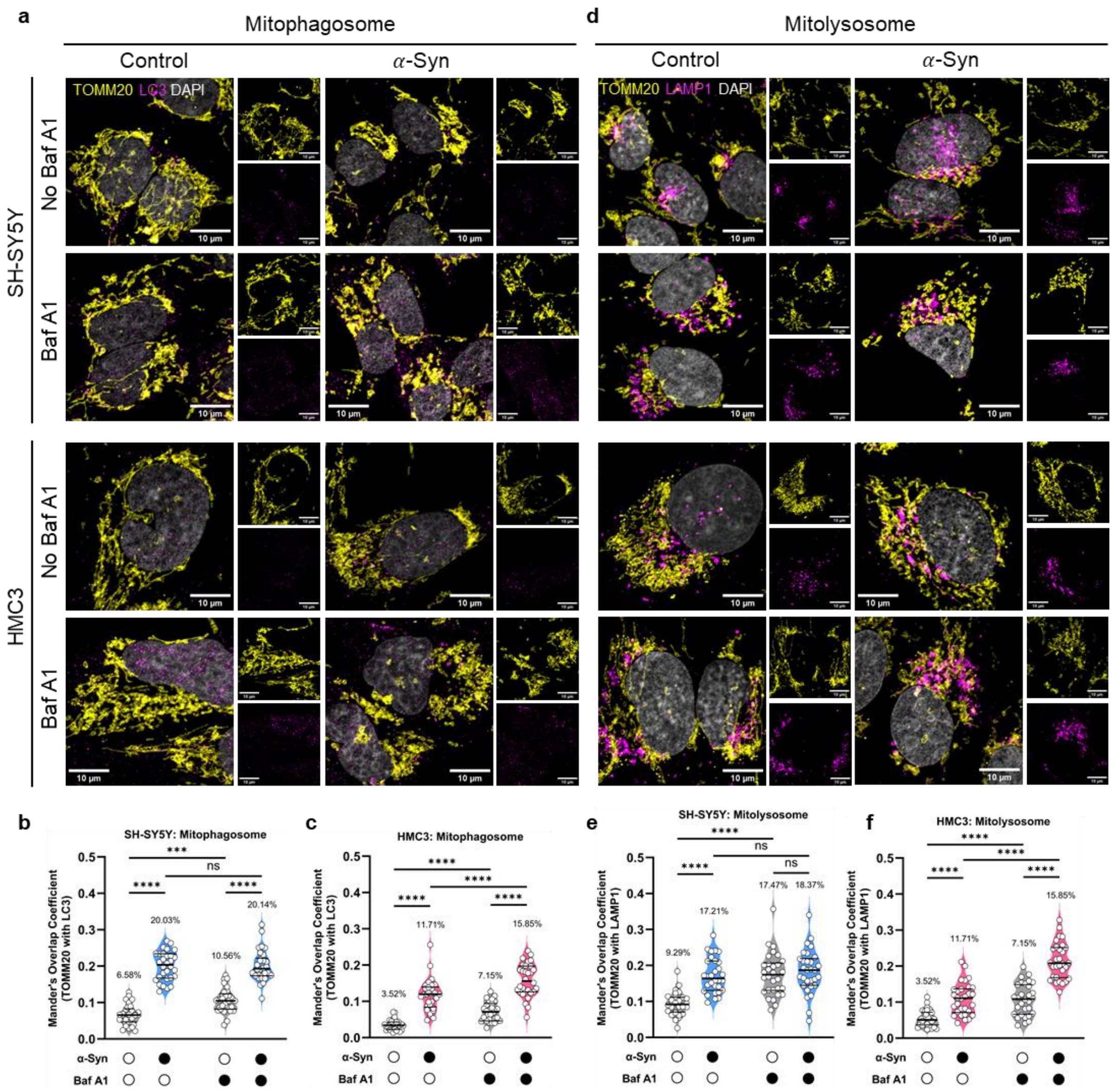


**Extended Data Fig. 2. Mitophagy flux in the presence of $\alpha$-Syn aggregates (related to Main Fig. 2).** (a) Representative confocal images of TOMM20+ mitochondria overlapping with LC3 (mitophagosomes) in different conditions in neuronal cells (top panels) and microglial cells (bottom panels). (b-c) Quantification of Mander's overlap coefficient of the fraction of TOMM20+ mitochondria overlapping with LC3 in neuronal cells (b) and microglial cells (c). N=3 independent experiments, n=30 cells per group. Statistical significance was analyzed using 2-Way ANOVA with Šídák's multiple comparison for (b), and uncorrected Fisher's LSD multiple comparison for (c). (d) Representative confocal images of TOMM20+ mitochondria overlapping with LAMP1 (mitolysosomes) in different conditions in neuronal cells (top panels) and microglial cells (bottom panels). (e-f) Quantification of Mander's overlap coefficient of the fraction of TOMM20+ mitochondria overlapping with LAMP1 in neuronal cells (e) and microglial cells (f). N=3 independent experiments, n=30 cells per group. Statistical significance was analyzed using 2-Way ANOVA with Šídák's multiple comparison for (e), and uncorrected Fisher's LSD multiple comparison for (f).

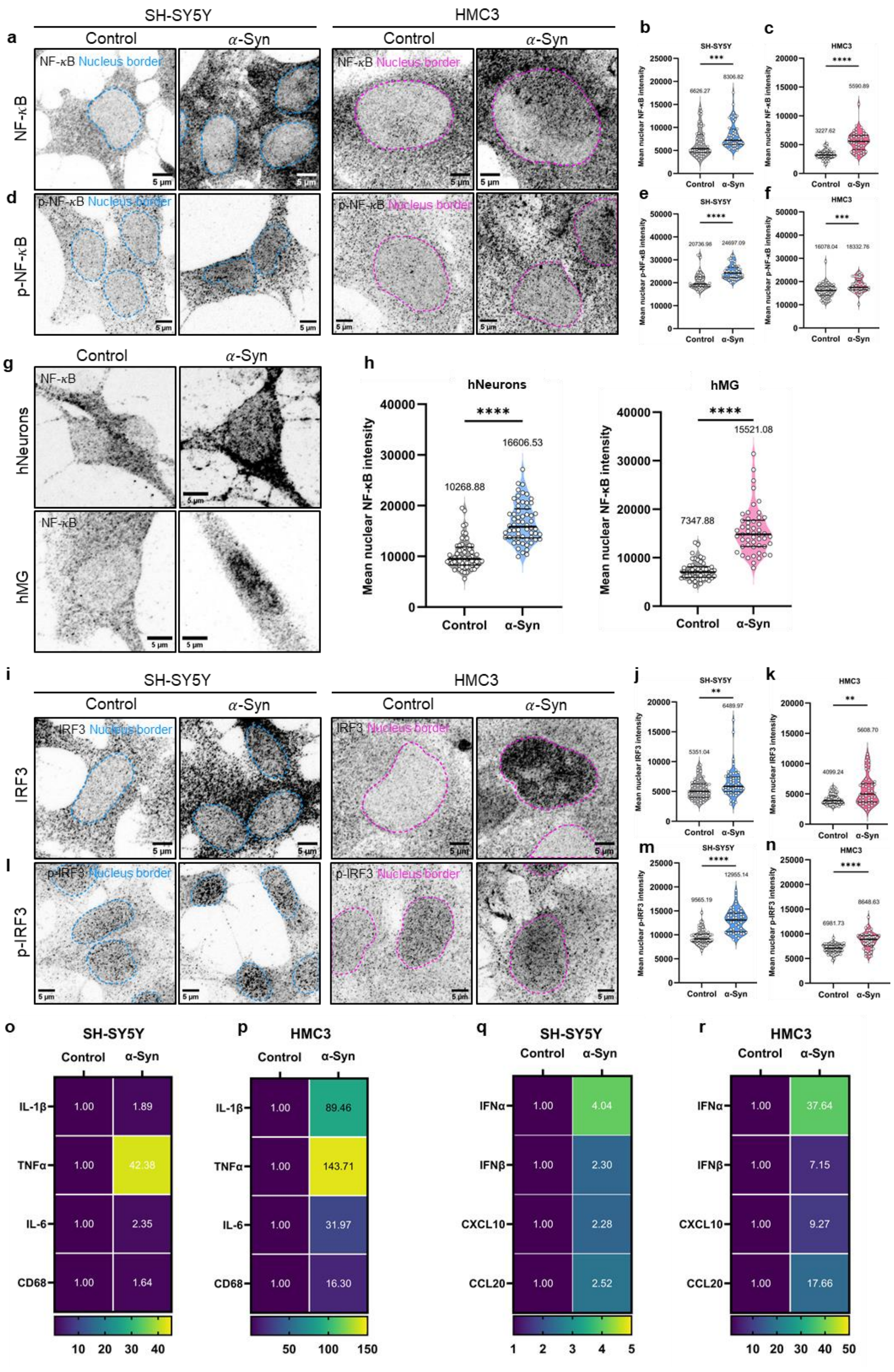


**Extended Data Fig. 3. Nuclear translocation of inflammatory factors and pro-inflammatory gene expression in response to $\alpha$-Syn aggregates (related to Main Fig. 4).** (a) Representative confocal images of neuronal cells (left panels) and microglial cells (right panels) to assess for nuclear levels of NF-$\kappa$B. (b-c) Quantification of mean nuclear NF-$\kappa$B intensity in neuronal cells (b) and microglial cells (c). N=3 independent experiments, n=50 cells per group. Statistical significance was analyzed using Mann-Whitney test. ***p<0.001, ****p<0.0001. (d) Representative confocal images of neuronal cells (left panels) and microglial cells (right panels) to assess for nuclear levels of p-Ser-536-NF-$\kappa$B. (e-f) Quantification of mean nuclear p-NF-$\kappa$B intensity in neuronal cells (e) and microglial cells (f). N=3 independent experiments, n=50 cells per group. Statistical significance was analyzed using Mann-Whitney test. ***p<0.001, ****p<0.0001. (g) Representative confocal images of hiPSC-derived neurons (hNeurons, left panels) and microglia (hMG, right panels) to assess for nuclear levels of NF-$\kappa$B. (h) Quantification of mean nuclear NF-$\kappa$B intensity in hNeurons and hMG. N=3 independent experiments, n=50 cells per group. Statistical significance was analyzed using Mann-Whitney test. ****p<0.0001. (i) Representative confocal images of neuronal cells (left panels) and microglial cells (right panels) to assess for nuclear levels of IRF3. (j-k) Quantification of mean nuclear IRF3 intensity in neuronal cells (j) and microglial cells (k). N=3 independent experiments, n=50 cells per group. Statistical significance was analyzed using Mann-Whitney test. **p<0.01. (l) Representative confocal images of neuronal cells (left panels) and microglial cells (right panels) to assess for nuclear levels of p-Ser-386-IRF3. (m-n) Quantification of mean nuclear p-IRF3 intensity in neuronal cells (m) and microglial cells (n). N=3 independent experiments, n=50 cells per group. Statistical significance was analyzed using Mann-Whitney test. ****p<0.0001. Data in all violin plots are represented as median and quartiles. Mean values are mentioned within the graphs. (o-r) Heatmaps of RT-PCR-based gene expression profiles of cytokines and type I interferons in neuronal cells (o and q) and microglial cells (p and r) in response to $\alpha$-Syn aggregates. N=3 independent experiments, fold changes for each gene (normalized to actin) relative to control are mentioned within the heatmaps.

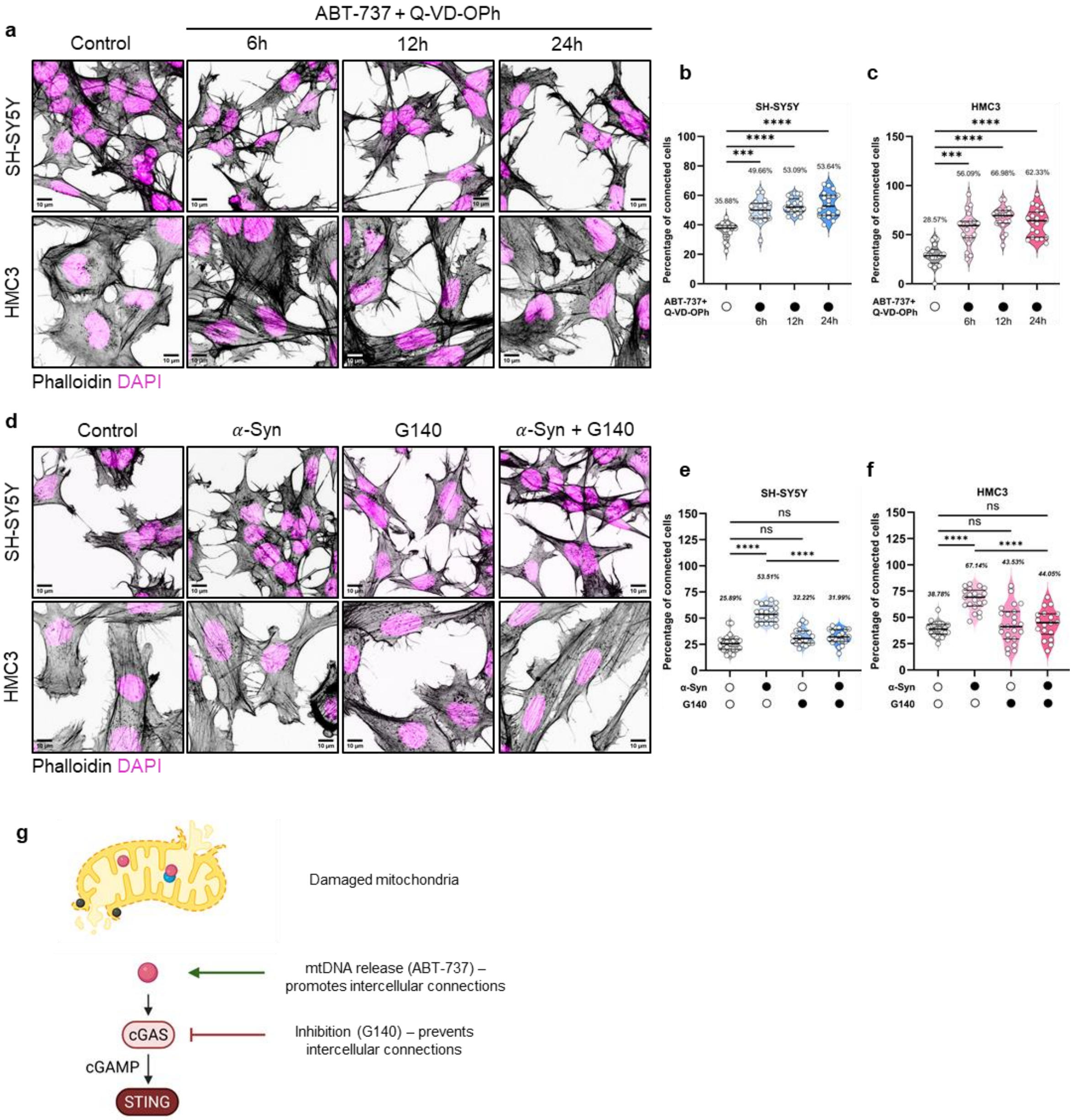


**Extended Data Fig. 4. Intercellular connections as a function of mtDNA release and cGAS activity (related to Main Fig. 5).** (a) Representative phalloidin-stained images of neuronal cells (top panels) and microglial cells (bottom panels) treated with limited MOMP-inducer ABT-737 (facilitating mtDNA release) and pan-caspase inhibitor Q-VD-OPh for the designated time points. (b-c) Quantification of the percentage of connected cells in different conditions for neuronal cells (b) and microglial cells (c). N=3 independent experiments, n=18-22 fields of views for neuronal cells, and 20 fields of views for microglial cells. Statistical significance was analyzed using Kruskal-Wallis test with Dunn's multiple comparison. ***p<0.001, ****p<0.0001. (d) Representative phalloidin-stained images of neuronal cells (top panels) and microglial cells (bottom panels) treated with $\alpha$-Syn alone, the cGAS inhibitor G140 alone, or co-treated with both. (e-f) Quantification of the percentage of connected cells in different conditions for neuronal cells (e) and microglial cells (f). N=3 independent experiments, n=21 fields of views for neuronal cells, and 18-23 fields of views for microglial cells. Statistical significance was analyzed using Brown-Forsythe and Welch ANOVA with Dunnett's T3 multiple comparison. ns: $p>0.05$, ****$p<0.0001$. Data represented as median and quartiles, with mean percentages of connections mentioned within the graphs. (g) Schematic representation of mtDNA release and cGAS as positive regulators of intercellular connections.

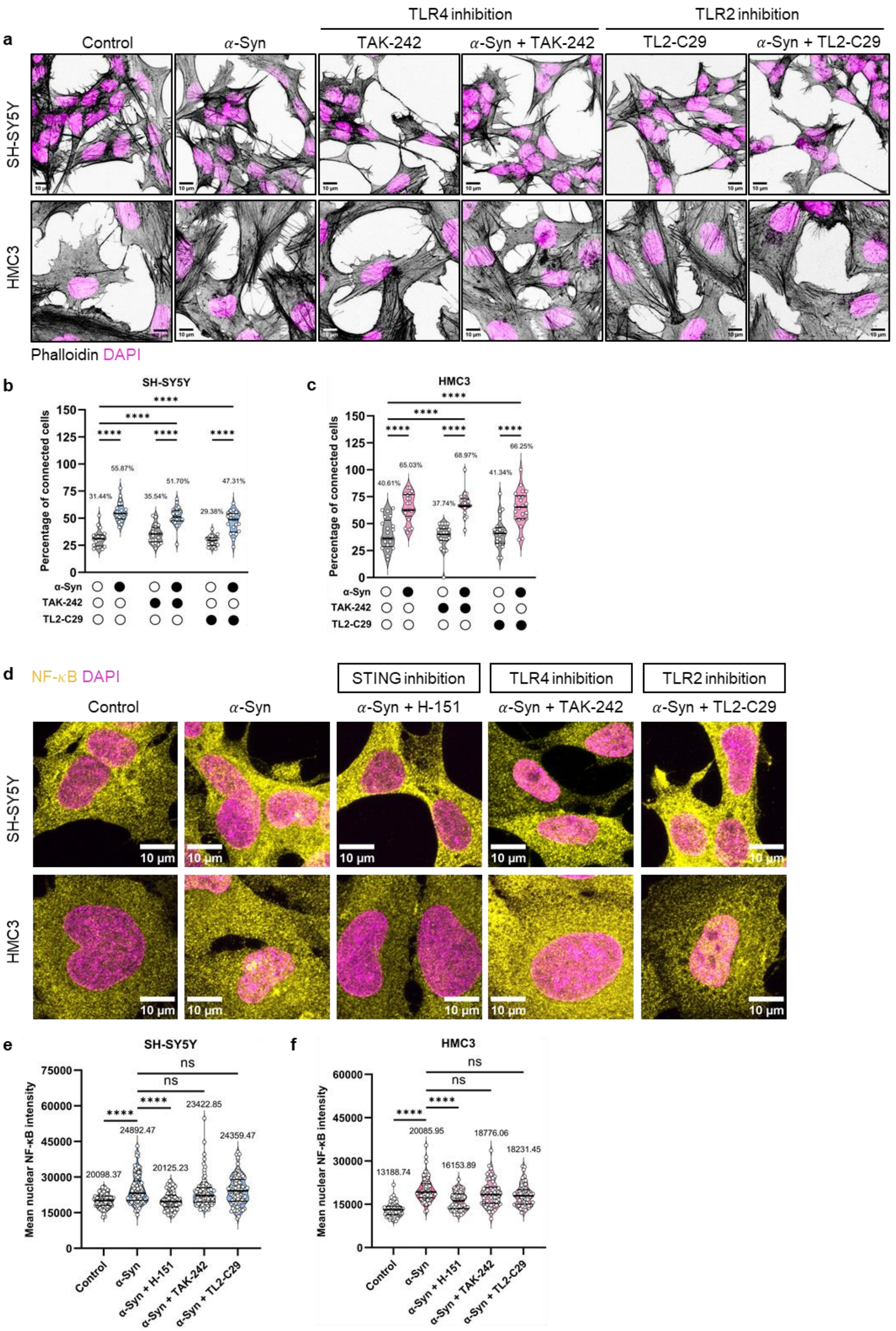


**Extended Data Fig. 5. Intercellular connections and NF-κB translocation to the nucleus in response to α-Syn after 16h of aggregate exposure is independent of TLR2/4.** (a) Representative phalloidin-stained images of neuronal cells (top panels) and microglial cells (bottom panels) in different conditions. (b-c) Quantification of the percentage of connected cells for neuronal cells (b) and microglial cells (c). N=3 independent experiments, n=20-22 fields of views for neuronal cells, and 21-22 fields of views for microglial cells. Statistical significance was analyzed using Two-Way ANOVA with Tukey's multiple comparison. ****$p<0.0001$. (d) Representative confocal images of neuronal cells (upper panels) and microglial cells (lower panels) to assess for nuclear translocation of NF-$\kappa$B upon $\alpha$-Syn exposure alone, or together with STING, TLR4, and TLR2 inhibitors. (e-f) Quantification of mean nuclear NF-$\kappa$B intensity in different conditions for neuronal cells (e) and microglial cells (f). N=3 independent experiments, n=100 cells per group for (e) and 60 cells per group for (f). Statistical significance was analyzed using Kruskal-Wallis test with Dunn's multiple comparison. ns: $p>0.05$, ****$p<0.0001$. Data represented as median and quartiles, mean values mentioned within the graphs.

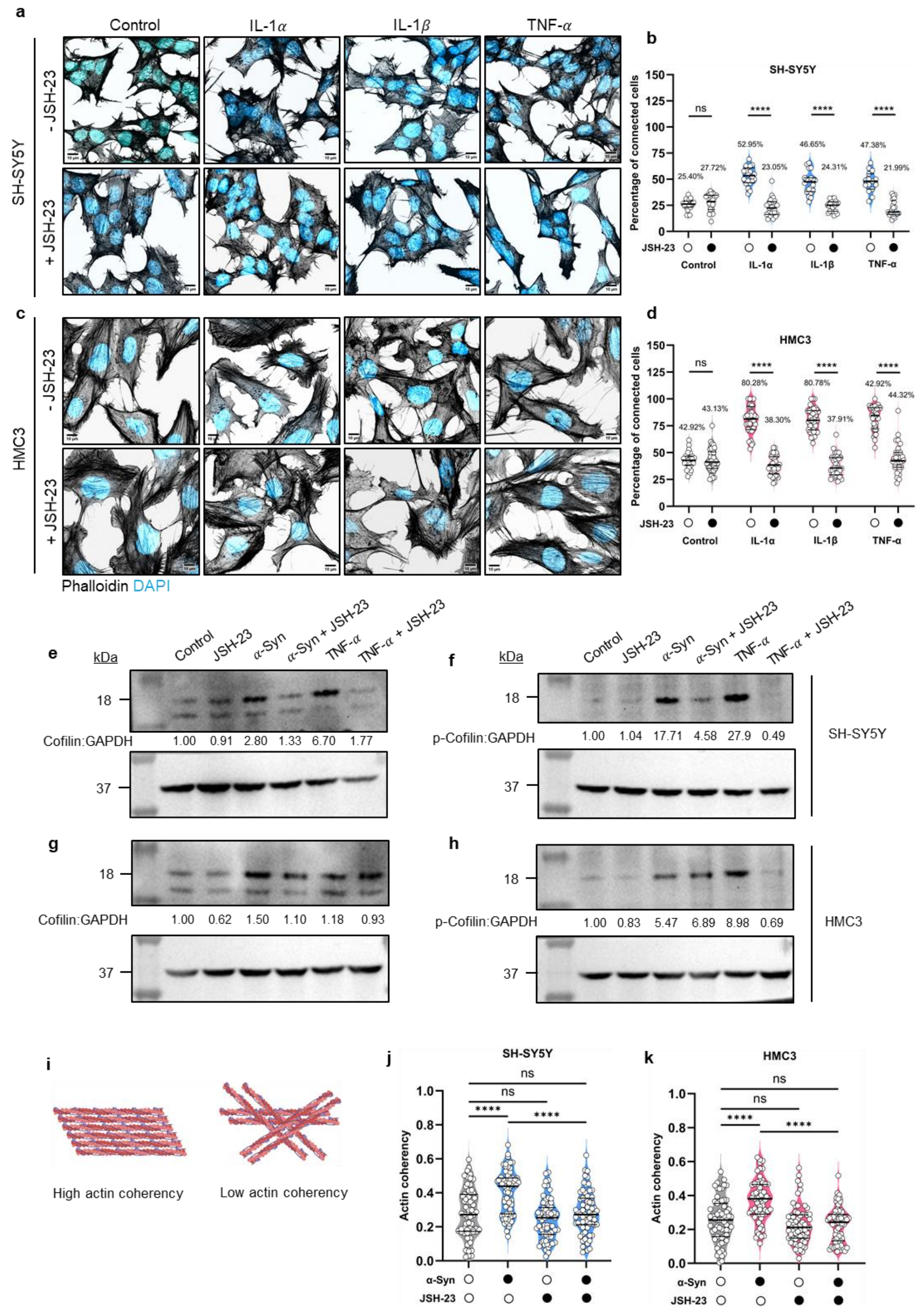


**Extended Data Fig. 6. Inflammation-induced intercellular connections are NF-$\kappa$B dependent, caused by changes in actin organization and Cofilin activity (related to Main Fig. 6).** (a) Representative phalloidin-stained images of neuronal cells treated with pro-inflammatory cytokines in the absence (top panels) or presence (bottom panels) of the NF-$\kappa$B inhibitor JSH-23. (b) Quantification of the percentage of connected neuronal cells in different conditions. (c) Representative phalloidin-stained images of microglial cells treated with pro-inflammatory cytokines in the absence (top panels) or presence (bottom panels) of the NF-$\kappa$B inhibitor JSH-23. (b) Quantification of the percentage of connected microglial cells in different conditions. For both (b) and (d): N=3 independent experiments, n=15-21 fields of views for neuronal cells and 23-38 fields of views for microglial cells. Statistical significance was analyzed using Two-Way ANOVA with Tukey's multiple comparison. ns: p>0.05, ****p<0.0001. Data represented as median and quartiles, with mean percentages mentioned within the graphs. (e-f) Representative western blots for Cofilin (e) and p-Ser3-Cofilin (f) in different treatment conditions for neuronal cells. N=2 independent experiments. Mean fold difference (normalized to GAPDH) relative to control is mentioned. (g-h) Representative western blots for Cofilin (g) and p-Ser3-Cofilin (h) in different treatment conditions for microglial cells. N=2 independent experiments. Mean fold difference (normalized to GAPDH) relative to control is mentioned. (i) Schematic representation of actin filament organization depicting high or low coherency. (j-k) Quantification of actin coherency for neuronal cells (j) and microglial cells (k) in control conditions, or treated with $\alpha$-Syn or JSH-23 alone, or co-treated with both. N=3 independent experiments, n=64-69 neuronal cells, and 63-66 microglial cells. Statistical significance was analyzed using Brown-Forsythe and Welch ANOVA with Dunnett's T3 multiple comparison. ns: p>0.05, ****p<0.0001. Data represented as median and quartiles.

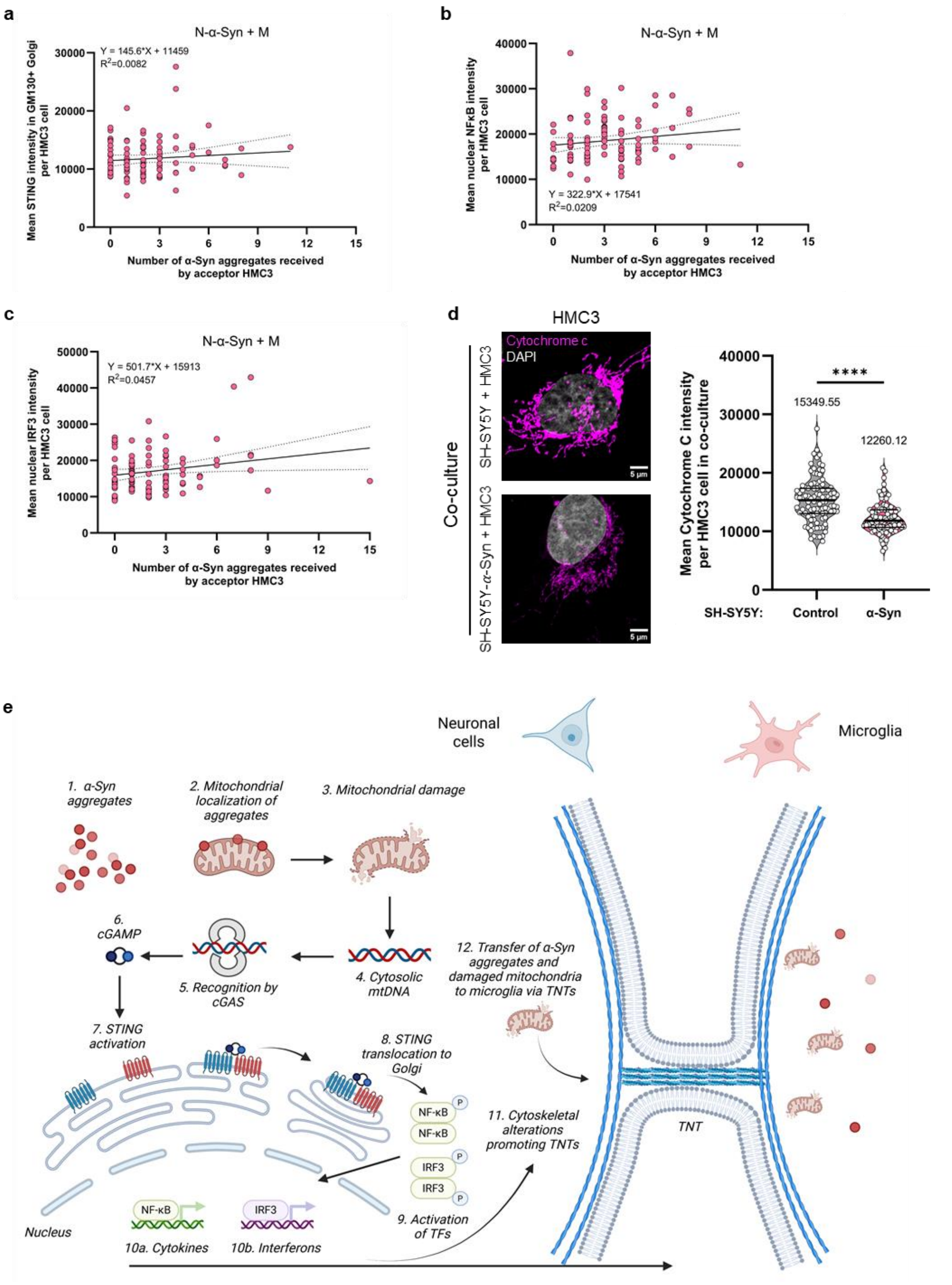


**Extended Data Fig. 7**. **Bystander inflammation in microglia is not correlated to the number of α-Syn aggregates they receive from neuronal cells (related to Main Fig. 8), and schematic depiction of the findings of this study**. (a-c) Correlation graphs of the number of $\alpha$-Syn aggregates received per cell, and STING translocation to GM130+ Golgi (a), NF-$\kappa$B translocation to the nucleus (b) and IRF3 translocation to the nucleus (c) in those very cells. Solid lines represent simple linear regression, and 95% confidence intervals are indicated by the dashed lines. (d) Representative confocal images and quantification of mean Cytochrome c intensity per microglial cell when in co-cultures with naïve (upper panel) or $\alpha$-Syn-burdened neuronal cells. N=3 independent experiments, n=100 microglial cells analyzed per group. Statistical significance was analyzed using Mann-Whitney test. ****$p<0.0001$. (e) Schematic representation of the main findings of this study.
